# Beyond Acoustic Cues: Olfactory-Mediated Avoidance of Bats by Crickets

**DOI:** 10.64898/2026.03.24.714006

**Authors:** Yannan Li, Wenhao Zhang, Jiaqi Wei, Hanhong Xu, Jiang Feng, Aiqing Lin

**Author notes:** For correspondence (AL); (JF).

## Abstract

The evolutionary arms race between insectivorous bats and their insect prey is a classic paradigm of acoustic predation and evasion, with insects having evolved sophisticated auditory countermeasures. Both bats and insects also rely heavily on olfaction for key behaviors, such as social communication. Moreover, predator-derived odors are well-established as risk cues in many other predator–prey systems. However, whether olfaction plays a role in the bat–insect arms race remains unknown. Here, we unveil a previously unknown olfactory dimension to this interaction. We demonstrated that the body odor of the insectivorous bat *Scotophilus kuhlii* triggered robust avoidance and electrophysiological antennal responses in a common cricket prey, *Loxoblemmus equestris*. We identified limonene as a behaviorally active volatile in bat odor that elicited electrophysiological responses in cricket antennae and was sufficient to elicit avoidance in crickets. Field experiments confirmed that limonene exposure reduced cricket calling activity, demonstrating the ecological relevance of this cue. Our findings establish that insects can detect and initiate avoidance of phylogenetically distant vertebrate predators via olfaction, a process that could be mediated by the elemental perception of individual odor compounds. This work broadens the sensory framework of a classic predator–prey system, and highlights olfactory eavesdropping as a functional strategy in phylogenetically distant predator–prey systems.

## Introduction

The evolutionary arms race between insectivorous bats and their insect prey represents a quintessential predator–prey system, characterized by the evolution of sophisticated acoustic countermeasures in insects (Conner and Corcoran, 2012; Ter Hofstede and Ratcliffe, 2016). As dominant nocturnal aerial insectivores, bats consume diverse taxa such as Lepidoptera, Coleoptera, and Diptera (Gong et al., 2025; Srilopan et al., 2025), providing vital biocontrol against agricultural pests like *Spodoptera frugiperda* and *Helicoverpa armigera* (Liu et al., 2024; Maine and Boyles, 2015). Over 50 million years of coevolutionary arms race, insects evolved sophisticated auditory countermeasures: ultrasound hearing, sound-absorbing scales, acoustic decoys, and ultrasonic jamming (Barber et al., 2022; Jacobs and Bastian, 2016; Zeng et al., 2011). Auditory detection, independently evolved in at least seven insect orders, is the most widespread adaptation (Ter Hofstede and Ratcliffe, 2016). Upon detecting bat echolocation calls, insects typically execute evasive flight maneuvers away from the predator source to reduce detection and predation risk (Jacobs and Bastian, 2016). However, the dominance of this acoustic narrative may have overshadowed other sensory channels (Stidsholt et al., 2025).

Notably, both bats and insects themselves rely heavily on olfaction for essential behaviors. Many bats (e.g., *Hipposideros armiger*, *Lasiurus cinereus*, *Phyllostomus hastatus*, and *Saccopteryx bilineata*) produce species-specific volatile organic compounds (VOCs) from specialized glands for social communication (Adams et al., 2018; Caspers et al., 2009; Stefaniak et al., 2025; Zhang et al., 2022). Conversely, insects possess exquisitely sensitive olfactory systems capable of detecting chemical cues (e.g., sex pheromones) over kilometers, critical for foraging, mate finding, and predator avoidance (Kannan et al., 2022). Thus, olfaction could, in principle, provide an additional sensory channel in the bat–insect dynamic. More broadly, predator-derived VOCs are well-established as risk cues that elicit adaptive avoidance behaviors in diverse predator–prey systems (Apfelbach et al., 2005, 2015; Ferrero et al., 2011; Rosen et al., 2015; Takahashi et al., 2005, 2008). Empirical studies demonstrate that prey organisms exhibit multifaceted antipredator responses, including enhanced vigilance, activity suppression, and abandonment of non-defensive behaviors (e.g., foraging), when exposed to predator odors (Cornelis et al., 2019; Hegab et al., 2015; Pérez-Gómez et al., 2015). Such chemically cued defenses are well-established within phylogenetic groups: vertebrate prey (e.g., rabbits, rodents, and ungulates) avoid mammalian predator odors (e.g., bobcats, foxes, and wolves), while invertebrate prey (e.g., various arthropods including crustaceans and insects) evade arthropod predators (e.g., aquatic insect larvae and spiders) (Kempraj et al., 2020; Monclús et al., 2005; Osada et al., 2014; Swihart et al., 1991; Weiss et al., 2018). These intraphyletic frameworks elucidate core principles of chemical communication and inform biological control strategies (Apfelbach et al., 2005; Dicke and Grostal, 2001).

However, a fundamental gap exists in understanding whether such olfactory eavesdropping can operate across the vast phylogenetic divide separating vertebrate predators and invertebrate prey (Apfelbach et al., 2015; Dicke and Grostal, 2001; Schoeppner and Relyea, 2005). Although olfactory interactions across broad taxonomic boundaries are widespread in nature, such as mosquitoes using host odors to blood-feed, elephants and moths sharing pheromonal components, and aroids chemically mimicking carrion to attract pollinating flies (Kang et al., 2023; Zaremska et al., 2022; Zhao et al., 2022), these interactions are primarily shaped by selective pressures tied to foraging, reproduction, or mutualisms, rather than by predation-related selection. Given the pronounced differences between vertebrates and invertebrates in olfactory receptor architectures, neural processing, and behavioral ecology (Kaupp, 2010; Wang et al., 2024), two pivotal and interconnected questions arise: First, can invertebrates detect and respond to volatile cues from phylogenetically distant vertebrate predators? Second, what is the mechanistic basis of such recognition? Specifically, does it require the holistic perception of complex, multi-component odor blends (configural processing), or can it be triggered by the detection of simple, elemental compounds (Apfelbach et al., 2015; Clifford and Riffell, 2013; Coureaud et al., 2022; Lei and Vickers, 2008)? Resolving these questions is essential to advancing theories of sensory evolution and chemical ecology beyond intraphyletic paradigms.

The bat–insect system presents an ideal model to address these questions. Despite the clear importance of olfaction to both taxa and its established role in predator–prey ecology, whether it plays any functional role in the iconic bat–insect arms race remains unexplored. In the present study, we hypothesized that insects might eavesdrop on bat VOCs as an early warning. Using the Asiatic lesser yellow house bat (*Scotophilus kuhlii*) and its cricket prey (*Loxoblemmus equestris*) as a model system, we tested whether bat body odor elicits avoidance in insects and if individual odor components were sufficient to trigger such responses. *S. kuhlii* (Mammalia: Chiroptera: Vespertilionidae) is widely distributed across Asian tropics and subtropics. In China, this species typically forms dense colonies in palm trees (*Livistona chinensis*). *S. kuhlii* preys on a variety of insects, including members of the orders Coleoptera, Orthoptera, and Lepidoptera (Zhu et al., 2012). This species forages on the wing over forests, farmlands, and grasslands, habitats shared with *L. equestris*. This spatial overlap creates predation opportunities, as the bats frequently hunt near the ground. In our study area, *L. equestris* dominates cricket assemblages in the foraging habitats of *S. kuhlii*, producing intense mating calls (primarily for mate attraction and territory defense) and frequently relocating by flight, behaviors that heighten its vulnerability to predators (Ren et al., 2023; Stevenson and Rillich, 2012). Notably, *S. kuhlii* emits a strong and distinctive odor. Following confirmation of this specific predator–prey relationship by DNA metabarcoding, we integrated behavioral, electrophysiological, chemical, and field evidence to demonstrate olfactory-mediated avoidance in *L. equestris* and identify its underlying mechanism.

## Results

### L. equestris is a prey species of S. kuhlii

To establish the predatory relationship between *S. kuhlii* and *L. equestris*, we conducted DNA metabarcoding analysis on fecal samples from 30 individual bats. Using mitochondrial 16S rRNA gene (16S) and cytochrome c oxidase subunit I gene (COI) markers, we obtained 426,832 insect sequence reads (323 operational taxonomic units, OTUs) and 1,805,577 insect sequence reads (534 OTUs), respectively. The diet of *S. kuhlii* was primarily composed of Coleoptera, Blattodea, Lepidoptera, Hemiptera, Orthoptera, Mantodea, Diptera, and Hymenoptera (Figure 1A). Within Orthoptera, Gryllidae was the dominant family, comprising 94% (16S) and 93.5% (COI) of Orthoptera sequences (Figure 1B), and was detected in 50% of bats (15/30 individuals). Phylogenetic analysis revealed at least seven cricket species among the prey (Figure 1C). Among these prey crickets, *L. equestris* was the most abundant species encountered at ground level in the bat foraging sites. This ground-level abundance was supported by insect surveys conducted over eight nights during fecal sample collection, which captured 80 crickets, 70 (87.5%) of which were *L. equestris* (Supplementary Table 1). Based on the crickets’ well-developed olfactory capacity, the confirmed predation by *S. kuhlii*, the high local abundance and accessibility of *L. equestris*, and its tractability in behavioral settings, we selected *L. equestris* for subsequent experiments testing olfactory responses to bat body odor.

**Figure 1.**
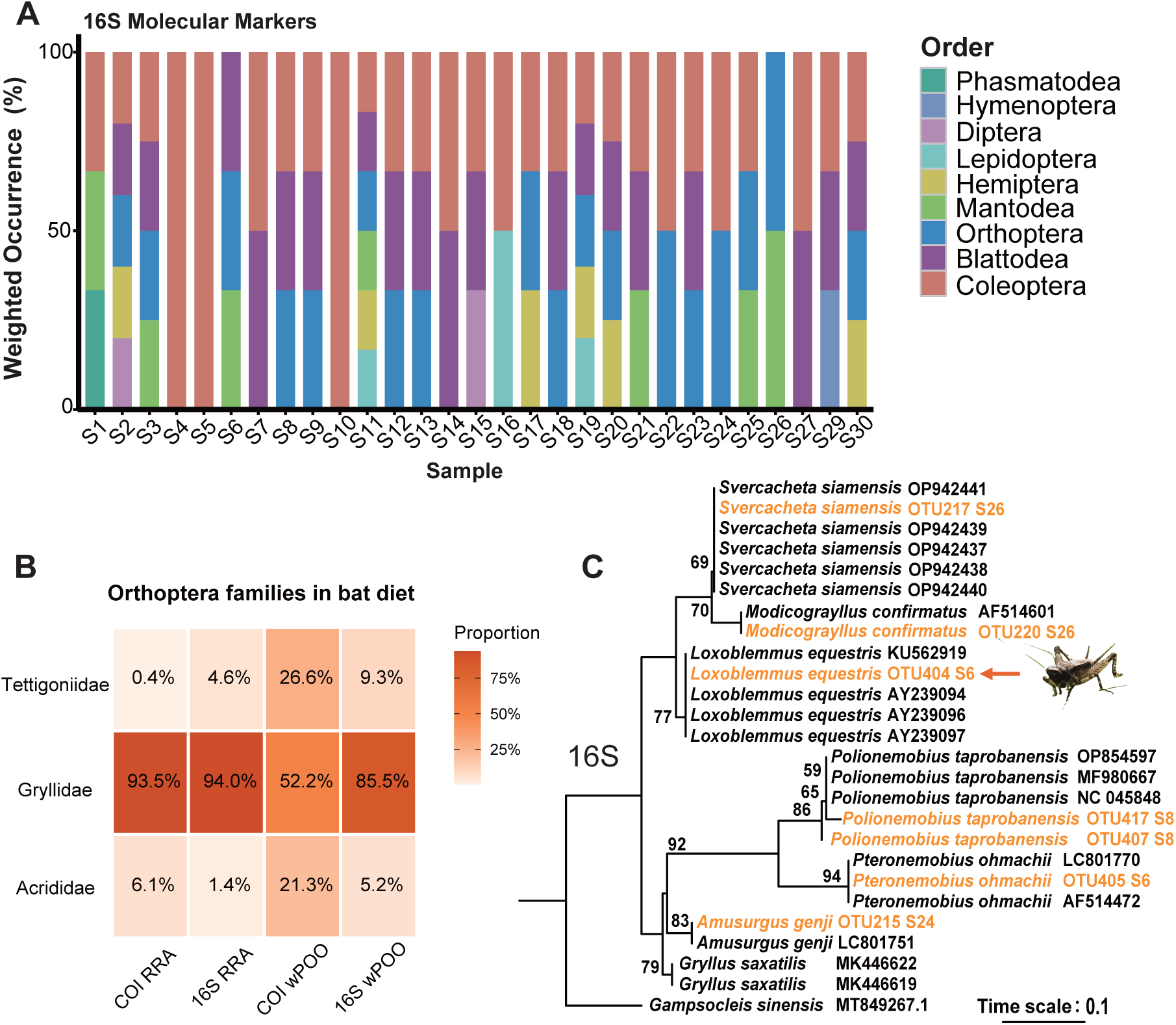
Dietary composition of *Scotophilus kuhlii*. (A) Order-level composition based on 16S gene sequences, analyzed using the weighted percentage of occurrence (wPOO). Individual bats are labeled S1–S30. (B) Proportions of Orthoptera families in the diet, detected using the COI and 16S genes and presented as both relative read abundance (RRA) and weighted percentage of occurrence (wPOO). Color intensity corresponds to proportion magnitude. (C) Maximum-likelihood phylogenetic tree of Gryllidae reconstructed from 16S sequences. Operational taxonomic units (OTUs) identified in this study are highlighted in orange; reference sequences from NCBI are shown in black. Associated bat individuals are listed after each corresponding scientific name. Bootstrap support values are indicated at the nodes.

### *S. kuhlii* body odor elicited behavioral avoidance and antennal responses in *L. equestris*

Behaviorally, in a two-choice olfactometer (Figure 2A), *L. equestris* strongly avoided the arm containing bat body odor. Upon release, 44 of 47 crickets (93.6%) chose the odor-free control arm, whereas only one entered the bat-odor arm and two shuttled without making a definitive choice (Chi-square test comparing avoiding vs. non-avoiding individuals: χ² = 35.76, *df* = 1, *P* < 0.001, φ = 0.87, 95% CI: 0.73 to 1.00; Figure 2B). Control trials with odor-free air in both arms showed no significant bias (13 of 24 vs. 11 of 24 crickets; Chi-square test, χ² = 0.17, *df* = 1, *P* = 0.68, φ = 0.08, 95% CI: 0 to 0.49; Figure 2B). Electrophysiological recordings using gas chromatography-electroantennographic detection (GC–EAD) confirmed that volatiles from *S. kuhlii* body odor are detected by *L. equestris* antennae. A mixture of bat odor volatiles from eight individuals elicited consistent antennal depolarizations (>0.1 mV) in all five crickets tested (Supplementary Figure 1). Parallel analysis by gas chromatography–mass spectrometry (GC–MS) identified six candidate compounds in the same odor samples. Among these, two compounds, namely 2,2-dimethylheptane and limonene, consistently co-eluted with EAD-active peaks (Figures 2C and 2D), identifying them as key antennal stimulants. No antennal responses were observed in odor-free controls tested on three crickets (Supplementary Figure 1). Together, these results establish that *L. equestris* perceives and avoids the body odor of its predator, *S. kuhlii*.

**Figure 2.**
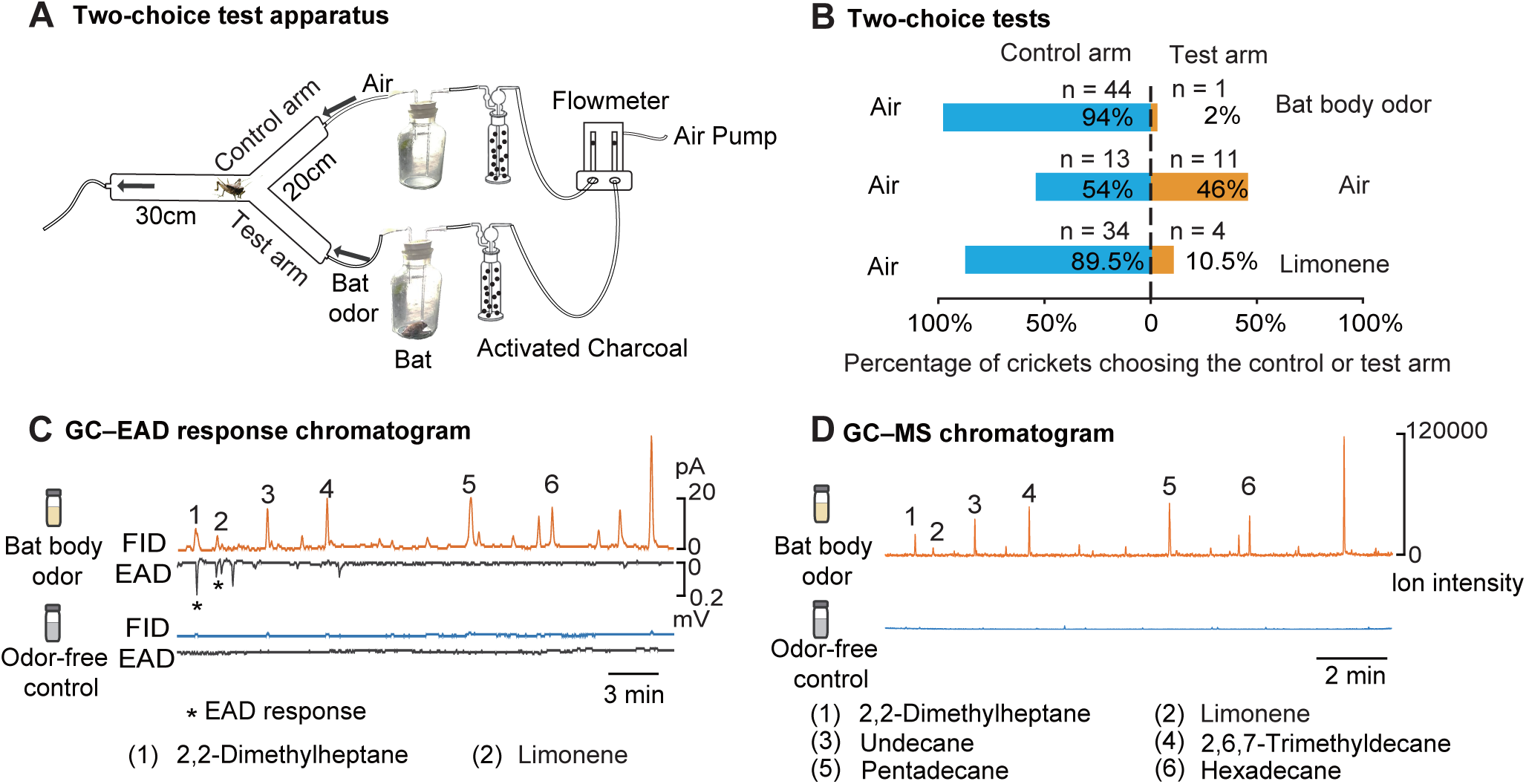
Behavioral and electrophysiological responses of *L. equestris* to *S. kuhlii* body odor. (A) Schematic of the Y-tube olfactometer used in two-choice assays. The cricket’s release position (junction) and airflow direction are indicated. (B) Choice percentages of crickets for the control versus test arm across three treatments: bat body odor, air without bat odor (control for odor bias), and limonene. (C) Representative gas chromatography–electroantennographic detection (GC–EAD) recordings. Upper panel: flame ionization detection (FID) chromatograms of bat body odor extract (brown) and an odor-free control (blue). Lower panel: corresponding EAD response (black) of a *L. equestris* antenna. Peaks 1–6 correspond to compounds identified by GC–MS (see panel D); asterisks mark compounds that elicited consistent antennal depolarizations (>0.1 mV). (D) Gas chromatography–mass spectrometry (GC–MS) total ion chromatograms of the same bat body odor sample (upper) and an odor-free control (lower). Identified compounds are labeled (peaks 1–6). Axes indicate retention time (x) and relative ion intensity (y).

### Snout secretions are the primary source of body odor in *S. kuhlii*

To identify the biological sources of bat body odor, we analyzed VOCs from hair, feces, and snout (pararhinal gland) secretions of nine bats using headspace solid–phase microextraction coupled with gas chromatography–mass spectrometry (HS–SPME–GC–MS). Snout secretions and hair shared similar hydrocarbon-rich VOC profiles, whereas fecal volatiles were distinct (Figure 3A). Individual-level terpenoid profiles further showed that limonene was detected in hair and snout secretion VOC collections but was absent from feces (Figure 3B and Supplementary Table 2). Principal component analysis revealed that the VOC profile of a pooled bat body odor sample (from eight bats) clustered more closely with those from snout secretions and hair than with feces (Figure 3C). As hair likely functions as a passive carrier, its surface VOCs probably originate from transferred snout secretions. These results point to snout secretions as the primary source of *S. kuhlii* body odor.

**Figure 3.**
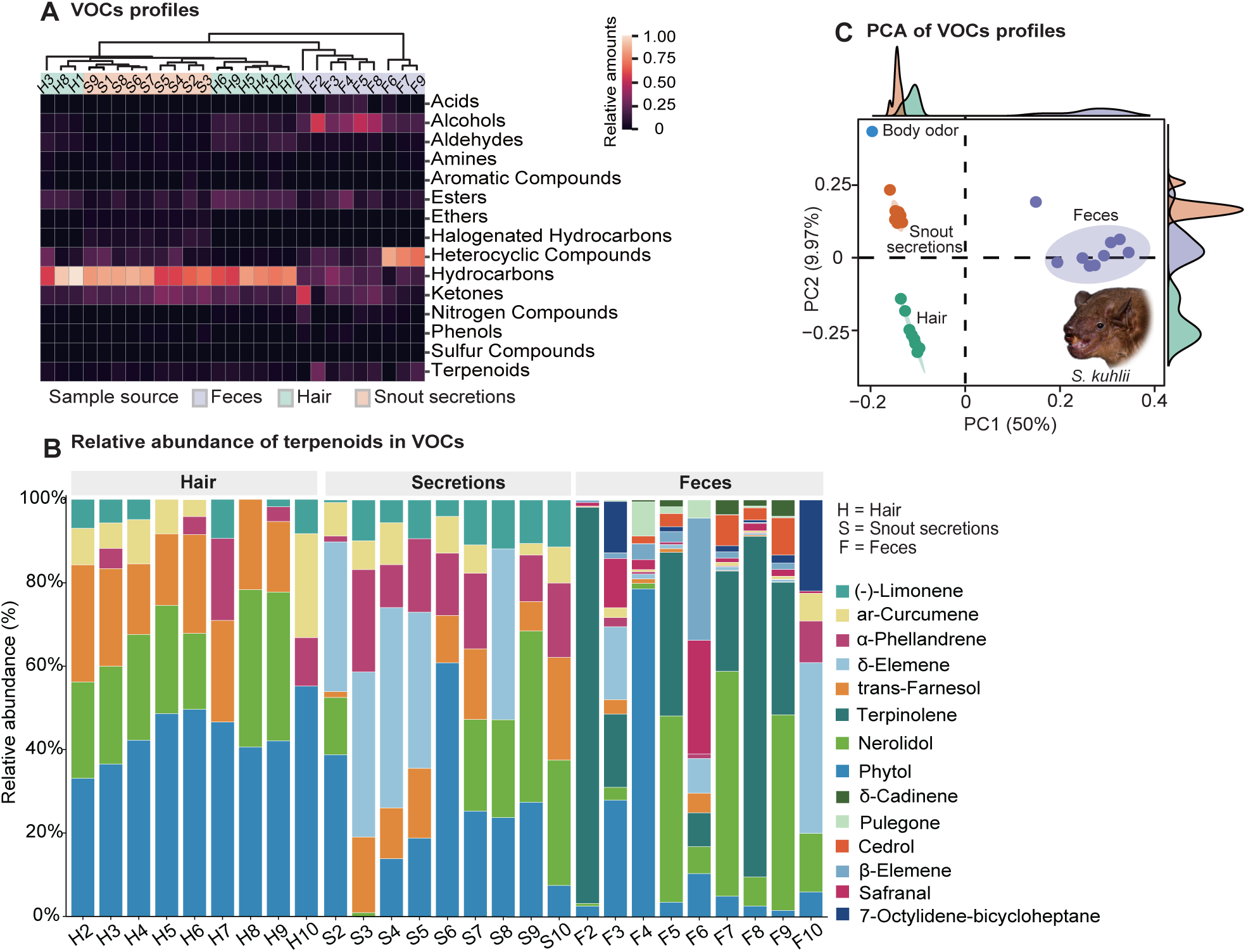
Volatile organic compound (VOC) profiles of potential odor sources in *S. kuhlii*. (A) Hierarchical clustering of VOC profiles from feces, hair, and snout secretions, based on compositional similarity (values normalized 0–1). All three odor sources were sampled from the same nine bats (27 samples total). (B) Individual-level relative abundance of terpenoid compounds in VOC collections from hair, snout secretions, and feces. Each stacked bar represents one VOC collection from an individual bat, and the segment corresponding to limonene indicates its presence and relative contribution in the sample. (C) Principal component analysis (PCA) of VOC profiles from the same samples, based on compound presence/absence data, with the addition of a pooled body odor composite (combined from eight bats) for comparison. Ellipses represent 90% confidence intervals, and marginal density plots are shown for each source.

### A single compound, limonene, triggers avoidance and reduces calling activity in *L. equestris*

To assess the behavioral relevance of identified volatiles, we measured antennal responses of *L. equestris* to four commercially available compounds previously detected in bat body odor: limonene, undecane, pentadecane, and hexadecane, using electroantennography (EAG). Compared to a hexane control, only limonene (10% v/v, 5.87 × 10^−7^ mol/µL) evoked a significant antennal response (repeated-measures ANOVA, *F*(4, 28) = 89.77, *P* < 0.001, η²*p* = 0.93; limonene vs. control: Bonferroni-corrected paired *t*-test, *t*(7) = 10.70, *P* < 0.001, Hedges’ *g* = 3.41; Figure 4A). EAG responses to limonene were concentration-dependent (repeated-measures ANOVA, *F*(5, 25) = 24.95, *P* < 0.001, η²*p* = 0.83; Figure 4B), with both 1% and 10% limonene solutions eliciting significantly stronger responses than the hexane control (Bonferroni-corrected paired *t*-tests, 1%: *t*(5) = 10.77, *P* < 0.001, Hedges’ *g* = 3.82; 10%: *t*(5) = 6.92, *P* = 0.005, Hedges’ *g* = 2.46).

**Figure 4.**
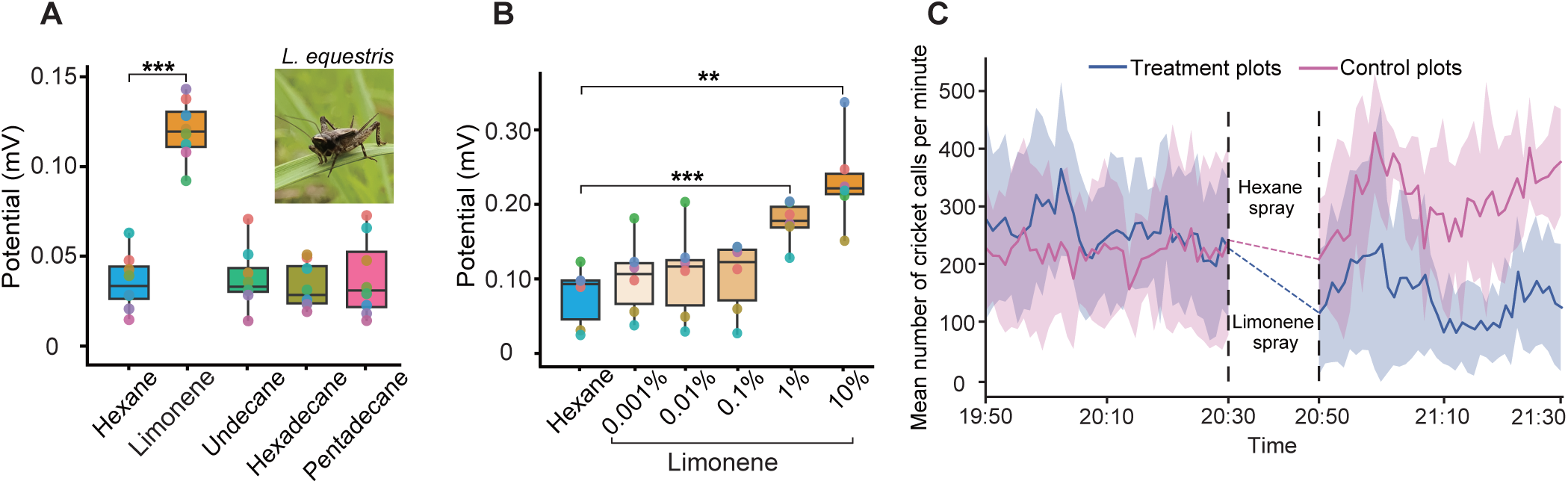
Electrophysiological and ecological responses of *L. equestris* to candidate bat body odor compounds. (A) Electroantennographic (EAG) responses to four candidate compounds (*n* = 8 individuals). Significance versus the hexane control is indicated (\*\**P* < 0.001; repeated-measures ANOVA with Bonferroni-corrected paired *t*-tests, see Results for exact *F*- and *P*-values and post-hoc comparisons). (B) Concentration-dependent EAG responses to limonene (*n* = 6 individuals). \*\**P* < 0.01, \*\*\**P* < 0.001 versus control (same statistical test, see Results). (C) Mean call rate over time (calls per minute) in field plots treated with limonene (*n* = 8 datasets) or a hexane control (*n* = 7 datasets). Shaded areas represent 95% confidence intervals (1,200 recordings total).

Notably, limonene alone was sufficient to elicit robust avoidance behavior in *L. equestris*. In a two-choice olfactometer assay, 34 of 38 *L. equestris* individuals (89.5%) chose the limonene-free control arm, demonstrating significant avoidance (Chi-square test, χ² = 23.68, *df* = 1, *P* < 0.001, φ = 0.79, 95% CI: 0.59 to 0.99; Figure 2B). Field experiments confirmed the ecological relevance of this avoidance. Prior to spray application during their peak nocturnal activity period, the mean call rate of *L. equestris* did not differ significantly between treatment and control plots (mean ± SD, 245 ± 143 vs. 207 ± 133 calls/min; GLMM, estimate = 0.10, SE = 0.25, 95% CI: −0.39 to 0.59, *Z* = 0.38, *P* = 0.720; Supplementary Table 3). When exposed to limonene, the mean call rate of *L. equestris* in treatment plots significantly decreased over the 40-min trial (post- vs. pre-exposure: mean ± SD, 130 ± 139 vs. 245 ± 143 calls/min; GLMM, estimate = −0.68, SE = 0.06, 95% CI: −0.80 to −0.57, *Z* = −11.90, *P* < 0.001; Figure 4C and Supplementary Table 3). In contrast, call rates increased significantly in control plots sprayed with hexane alone (post- vs. pre-exposure: 298 ± 126 vs. 207 ± 133 calls/min; GLMM, estimate = 0.40, SE = 0.06, 95% CI: 0.28 to 0.51, *Z* = 6.68, *P* < 0.001; Figure 4C and Supplementary Table 3), a pattern consistent with their natural crepuscular activity peak. Together, these results demonstrate that limonene, a single component of bat odor, is sufficient to induce avoidance and reduce call activity in *L. equestris* under natural conditions.

## Discussion

Our study provides direct experimental evidence that an insect species can detect and avoid a phylogenetically distant vertebrate predator via olfaction. We demonstrate that the body odor of the insectivorous bat *S. kuhlii* triggers robust avoidance and electrophysiological responses in the cricket *L. equestris*, and that a single compound, limonene, is sufficient to elicit this avoidance in the laboratory and suppress calling in the field. These findings fundamentally expand the sensory narrative of a classic predator–prey system and offer novel insights into the mechanisms of olfactory detection and chemical eavesdropping in interactions across broad phylogenetic divides.

### Olfaction as a novel and functional channel in bat–insect interactions

For decades, the bat–insect arms race has served as a paradigm for acoustic coevolution, with insect auditory adaptations to echolocation being meticulously documented (Conner and Corcoran, 2012; Ter Hofstede and Ratcliffe, 2016). This dominant narrative, however, may have inadvertently narrowed the research focus, contributing to relatively less exploration of alternative sensory channels beyond audition (Stidsholt et al., 2025). Many bats produce body odor in the form of VOCs from specialized glands for intraspecific communication (Adams et al., 2018; Caspers et al., 2009; Stefaniak et al., 2025; Zhang et al., 2022). Here, we found that the body odor of *S. kuhlii*, which originates primarily from snout secretions, elicits robust avoidance and antennal responses in the cricket *L. equestris*. This suggests that insects may eavesdrop on predators’ intraspecific olfactory signals to mitigate predation threat. Our discovery thus reveals a previously overlooked olfactory dimension to this classic predator–prey interaction. This finding aligns with the established importance of multimodal predator detection, challenging the presumed supremacy of acoustics in this particular system (Jacobs and Bastian, 2016).

While our laboratory assays demonstrate clear olfactory-mediated avoidance in crickets, the ecological dynamics of a direct bat–insect encounter in the wild are more complex. Nonetheless, our field experiment provides critical, ecologically relevant support for the anti-predator function of this cue. The significant reduction in cricket calling activity upon limonene exposure—whether due to dispersal from the limonene-treated plots or direct cessation of signaling—constitutes a functionally adaptive anti-predator response, as both outcomes would reduce acoustic attraction and lower predation risk (Belwood and Morris, 1987; Hedrick and Kortet, 2006). This behavioral response confirms the ecological relevance of the olfactory cue, consistent with the evolutionary logic of eavesdropping on predator signals (Hermann and Thaler, 2014).

The functional advantages of this olfactory channel likely complement, rather than replace, established auditory defenses. The high sensitivity and long-range nature of insect olfaction may facilitate earlier predator detection, particularly against bats employing low-intensity or ‘stealth’ echolocation strategies that challenge acoustic detection alone (Corcoran and Conner, 2017; Goerlitz et al., 2010). More broadly, it could provide a vital, generalized defense for insect taxa lacking specialized ultrasonic hearing, including many beetles and non-hearing moths that are common bat prey (Göpfert and Hennig, 2016; Lin et al., 2023). The suppression of calling activity we observed exemplifies a key benefit: reducing acoustic conspicuousness. Insect calls can serve as a potential cue for certain bats, either through passive hearing or by enhancing the prey’s detectability within the returning echo (Alem et al., 2011; Falk et al., 2015; Prakash et al., 2021). We propose that olfaction in this system acts as a general alert mechanism, priming vigilance and enabling more informed deployment of evasive maneuvers triggered by closer-range acoustic (including echolocation) cues. In this integrated model, olfactory and auditory channels might operate across complementary spatial and temporal scales, forming a more robust, layered sensory defense system. Ultimately, direct observations of bat–insect encounters within manipulated olfactory environments will be crucial to fully quantify the survival benefit conferred by this eavesdropping channel under real predation pressure.

### Implications for cross-phylum sensory ecology and the mechanism of eavesdropping

Beyond the specific bat–insect model, our work addresses a central question in sensory ecology: how chemical eavesdropping operates within predator–prey systems between phylogenetically distant taxa with fundamentally divergent olfactory systems (Adams et al., 2020; Emerson and Johnson, 2024; Kaupp, 2010). While intraphyletic kairomone detection is well-established (e.g., rodents avoiding carnivore odors, aphids responding to ladybug chemicals), compelling experimental evidence for such olfaction-mediated recognition across broad phylogenetic divides has been limited (Apfelbach et al., 2005; Ferrari et al., 2007; Tanis et al., 2018). By linking physiological detection to behavioral avoidance within a confirmed predator–prey relationship, we demonstrate that functional cross-phylum olfactory eavesdropping is attainable, proving this capacity persists despite profound divergence in underlying olfactory circuitry.

A key question concerns the perceptual mechanism underlying such olfactory eavesdropping: does it require a complex, species-specific odor blend (“configural processing”), or can it be initiated by simple, “elemental” compounds (Apfelbach et al., 2015; Coureaud et al., 2022; Lei and Vickers, 2008)? Our identification of limonene as a single, sufficient elicitor of avoidance supports an elemental perception strategy in this cross-phylum context. This mechanism is evolutionarily parsimonious. Evolving high-sensitivity receptors for a reliable kairomone bypasses the need to decode complex vertebrate odor profiles (Schoeppner and Relyea, 2005), likely being more efficient than evolving sophisticated neural circuits for configural processing (Haverkamp et al., 2018; Niven and Laughlin, 2008). Functionally, elemental perception enables a rapid, high-sensitivity response to trace amounts (Stengl, 2010), potentially allowing prey to identify threats at lower concentrations or greater distances than required for complex blends (Cardé and Willis, 2008; Minor and Kaissling, 2003). Thus, elemental perception emerges as a plausible and efficient strategy for initiating anti-predator behavior against phylogenetically remote threats.

Although limonene reliably induced avoidance behaviour in crickets, two related questions still merit careful consideration. One question is whether the limonene we identified genuinely originates from bats or reflects contamination during sampling. Limonene is common in plants and numerous consumer products (Boncan et al., 2020; Schuman, 2023), making its endogenous production by a mammal seem unusual. Nevertheless, multiple lines of evidence militate against contamination. First, we adhered to rigorous protocols. For example, all instruments were cleaned with ethanol and oven-dried before each use; bats were housed in stainless-steel cages, and cloth bags had been rinsed with purified water. Second, limonene was absent from all blank controls, including empty-chamber air samples and clean swabs, and was not detected in bat fecal samples. In contrast, it was consistently identified in hair and snout-secretion samples from bats. Third, independent studies have similarly identified limonene in the secretions of other bat species (Faulkes et al., 2019; Zhang et al., 2022). Furthermore, emerging evidence indicates skin-associated microbes may contribute to bat volatile profiles, with some taxa possessing enzymes involved in terpene biosynthesis (Sun et al., 2026). Taken together, these observations point towards an endogenous or microbe-mediated source, although the exact biosynthetic pathway remains to be determined.

The second question is how crickets might distinguish bat-derived limonene from environmental sources of this compound, given its prevalence in mint, citrus peel, pine and other non-threatening plants (Boncan et al., 2020; Schuman, 2023). It seems implausible that crickets could simply rely on limonene per se to differentiate a bat from a leaf. Two non-exclusive mechanisms could help resolve this issue. First, limonene need not be the only olfactory cue mediating risk perception. Our findings establish the sufficiency of limonene as an avoidance trigger, but do not preclude a role for other odor components. The crickets’ antennal responses to other bat volatiles in our GC–EAD analyses suggest more complex peripheral perception.

Additional compounds, either alone or in synergistic blends, may modulate the full behavioral response in nature. Therefore, elemental perception via limonene likely represents one effective strategy within a broader olfactory toolkit available to insects. Second, crickets may discriminate bat-derived limonene through context-specific cues (e.g., temporal and spatial patterning, co-occurrence with other bat-specific compounds) to minimize false alarms. Comparative studies on enantiomeric specificity and detection thresholds of cricket olfactory sensory neurons will be essential. Equally critical will be future efforts to quantify natural bat odor composition, limonene release rates, ambient exposure concentrations, and odor-plume dynamics, which together will inform ecologically valid stimulus design in controlled assays. Critically, our field data confirm that limonene exposure in nature robustly triggers an adaptive anti-predator response, irrespective of the precise discrimination mechanism.

In conclusion, our integrated approach reveals that the body odor of *S. kuhlii* triggers avoidance in *L. equestris* and that a single component, limonene, is sufficient to elicit this behavior in the laboratory and suppress calling in the field. We have uncovered a novel olfactory axis in the classic bat–insect arms race, demonstrating that chemical cues constitute a functional sensory channel for predator detection. More broadly, it provides evidence that olfactory eavesdropping can operate across vast phylogenetic distances within predator–prey systems. Elemental perception of individual odor compounds could represent a parsimonious strategy for such cross-phylum threat detection. Future work on the neural circuitry, phylogenetic distribution, and ecological contingencies of this capability will elucidate how such olfactory-mediated eavesdropping evolves and operates across the tree of life. The identification of behaviorally active volatiles like limonene may offer a starting point for the bioinspired development of novel insect repellents (Rasmann et al., 2005), but such applications must be informed by a deeper understanding of the ecological context in which these cues are perceived and interpreted.

## Materials and methods

### Diet composition and prey availability of *Scotophilus kuhlii*

To confirm the predator–prey relationship between *S. kuhlii* and *L. equestris*, we analyzed the diet of *S. kuhlii* via DNA metabarcoding of fecal samples. On July 10, 2022, we captured 30 individuals of *S. kuhlii* (21 females and nine males) from a roosting colony in a Chinese fan palm in Guangzhou, China, using hand nets. Sampling occurred during the bats’ initial return to the roost following nightly foraging activity, a period associated with high insect consumption. Each bat was placed individually in a thoroughly washed cotton bag, and fecal samples were collected after two hours. The samples were immediately transferred to 2-mL Eppendorf tubes (Corning, Cat# 430659) and stored at –20 °C. Bats were released unharmed after sample collection.

From each fecal sample, a 100-mg subsample was taken for DNA extraction using an Omega Bio-tek DNA extraction kit according to the manufacturer’s instructions. We targeted a 225-bp fragment of the mitochondrial cytochrome *c* oxidase subunit I (COI) gene with primers LCO-1490 (5′-GGTCAACAAATCATAAAGATATTGG-3′) and ZBJ-ArtR2c (5′-WACTAATCAATTWCCAAATCCTCC-3′), and a 105-bp fragment of the mitochondrial 16S ribosomal DNA with primers Coleop_16Sc (5′-TGCAAAGGTAGCATAATMATTAG-3′) and Coleop_16Sd (5′-TCCATAGGGTCTTCTCGTC-3′) (Lin et al., 2023). Polymerase chain reaction conditions followed published protocols (Lin et al., 2023). Amplified products of all 64 samples (60 from bat feces and four extraction blanks as negative controls) were sequenced on an Illumina MiSeq PE300 platform following the standard operating procedures at Majorbio Bio-Pharm Technology Co., Ltd. Ultimately, using the 16S molecular marker, we obtained insect sequences from 29 bats; no insect sequences were detected for one individual (bat identity: S28). Using the COI molecular marker, we obtained dietary sequences from 30 individual bats.

Dietary analysis was performed following established bioinformatic protocols (Chang et al., 2019; Lin et al., 2023). Briefly, after quality filtering and merging paired-end reads, DNA sequences were clustered into Operational Taxonomic Units (OTUs) at a 97% similarity threshold using Usearch v11.0 (Edgar, 2016). Taxonomic assignment was conducted by comparing representative sequences from each OTU against the GenBank and BOLD databases, based on the following criteria: (a) sequences with >98% identity to a single species were assigned to that species; (b) those matching multiple congeners within the same genus (>98%) were assigned to the genus level; and (c) sequences matching multiple genera within a family or only family-level references were assigned to the family level. No contaminant insect sequences were detected in the negative controls. Only insect sequences were retained for diet analysis. We quantified bat diet composition using the relative read abundance (RRA) and the weighted percentage of occurrence (wPOO) of prey orders, families and species (Deagle et al., 2018). For RRA, total read counts were normalized to 80,000 reads per sample. The wPOO was calculated from occurrence frequency tables standardized to a total occurrence frequency of 100 across all samples.

To achieve species-level resolution within Gryllidae (crickets), we implemented a phylogenetic approach (Ross et al., 2008). Representative sequences of each Gryllidae OTU were aligned with their top five BLAST matches from NCBI, and duplicate sequences were removed. Multiple sequence alignment was performed in MEGA v7.0 (Kumar et al., 2016), and a maximum-likelihood (ML) phylogeny was reconstructed using IQ-TREE v1.6.12 under the best-fit substitution model (16S: TPM3u+F+G4; COI: TIM2+F+I+G4) (Nguyen et al., 2015), with *Gampsocleis sinensis* sequences as the outgroup. OTUs clustering within established species-level clades were assigned to the corresponding species.

To assess prey availability, we surveyed insects in a grassland foraging site of *S. kuhlii* approximately one kilometer from its roost. Sampling was carried out on eight nights between July and September 2022 (19:30–21:30) using one ultraviolet light trap (BY201, Beijing Baoyuan Xingye Technology Co., Ltd., China) and one Malaise trap (BY201, Beijing Baoyuan Xingye Technology Co., Ltd., China). Lepidoptera (moths) and Gryllidae were the most abundant taxa collected. Preliminary Y-tube olfactometer trials indicated that moths exhibited minimal movement in the Y-tube, precluding reliable assessment of odor-mediated behavior, whereas crickets showed consistent locomotor activity. We therefore focused on estimating the relative abundance of Gryllidae species. *L. equestris* was selected for subsequent olfactory experiments due to its high local abundance and confirmed status as a natural prey item of *S. kuhlii* (see Results).

### Behavioral and electrophysiological assays of cricket responses to *S. kuhlii* body odor

We employed two-choice tests in a Y-tube olfactometer (Shelai Instrument Platform, China) to assess whether the body odor of *S. kuhlii* elicits avoidance behavior in the cricket *L. equestris*. Individuals of *L. equestris* were collected nightly (19:30–21:30) from the bat’s foraging habitat (see the section above) and acclimated in insect cages with water provided in a temporary laboratory until testing.

The olfactometer consisted of a Y-shaped glass tube (inner diameter: 35 mm; main arm length: 200 mm; side arm length: 200 mm). One side arm was connected in series to a glass chamber housing one of eight adult *S. kuhlii* individuals (randomly selected for each trial) and a charcoal filter bottle; the other side arm was connected to an empty glass chamber and a charcoal filter bottle. Both charcoal bottles contained activated carbon for air purification. An air pump supplied airflow, regulated to 1 L/min in each arm by flowmeters.

Behavioral trials commenced at 23:00 under complete darkness. Before each trial, the Y-tube was cleaned with ethanol and oven-dried. Odorless sponges (diameter: 35 mm, thickness: 15 mm; Wuhan Hemida Technology Co., Ltd., China) were placed at the end of each arm to prevent the crickets from escaping. After confirming system airtightness, a single cricket was introduced at the junction (center) of the Y-tube, and its behavior was recorded using an infrared video camera (4K Ultra; Sony, China). Preliminary observations confirmed that crickets typically initiated exploration of the apparatus arms within three seconds of being introduced. Therefore, video recording began once a cricket started exploring the two side arms. A cricket was scored as “avoiding” the bat odor if it entered the control arm and remained there continuously for 10 s. It was scored as “not avoiding” the bat odor if it entered the test arm and remained there continuously for 10 s. Individuals that moved back and forth between the two arms without remaining in either arm for at least 10 s were also classified as “not avoiding”. We selected a 10-second duration as the criterion because the crickets were highly mobile, and this period was sufficient for them to fully explore the olfactometer. A total of 47 crickets were tested, with each individual used in a single trial. The left/right position of the bat odor source was randomized between trials to control for side bias. A separate control experiment with 24 crickets was performed under identical conditions, except that no bat was placed in the glass chamber. Behavioral responses were recorded following the same procedure. To minimize observer bias, the video recordings were analyzed by an experimenter who was blinded to the treatment conditions.

We collected the body odor of *S. kuhlii* (i.e., its volatile organic compounds, VOCs) for chemical characterization and electrophysiological bioassay using gas chromatography–mass spectrometry (GC–MS) and gas chromatograph–electroantennographic detection (GC–EAD) with cricket antennae, respectively. Samples were collected from eight adult individuals (four females and four males) using a dynamic headspace sampling system equipped with Porapak-Q adsorbent columns. We selected Porapak-Q because bat body odor is rich in hydrocarbons, which are non-polar or only weakly polar, and Porapak-Q has a high adsorption capacity for these compounds. The system consisted of a vacuum pump (QC-1S; Beijing Institute of Labour Protection Science, China), a charcoal filter for air purification, a 5 L glass chamber housing the bats, and a Porapak-Q column (glass tube, 4 mm internal diameter, 200 mg of Porapak-Q 80/100 mesh; Supelco, USA) (Wang et al., 2023). Prior to sampling, all glassware was thoroughly rinsed with ethanol and dried in an oven at 120°C, and volatile odor collection was conducted in a dedicated odor-free room to minimize environmental contamination. Each adsorbent column was cleaned by rinsing sequentially with 3 mL of methanol and dichloromethane (DCM), followed by thermal conditioning under nitrogen at 180 °C for 30 minutes. During VOC collection, purified air from the pump was passed through the system at a flow rate of 500 mL/min for two hours. Control samples were collected from empty glass chambers following the same procedure. After sampling, VOCs were eluted from each column with 1 mL of DCM, and the extracts were concentrated under a gentle nitrogen stream to a final volume of 1 mL. All samples were stored at –20 °C until subsequent analysis by GC–EAD and GC–MS. The eight *S. kuhlii* used for odor collection and behavioral assays were captured from a roosting palm tree on 1 October, 2023. Bats were housed socially in cages (50 × 50 × 50 cm) in a temporary laboratory maintained at 28 °C and 60% relative humidity. Mealworm larvae (*Tenebrio molitor*) and water were provided daily following nightly experiments.

The antennae are a primary sensory organ for detecting both chemical and mechanical cues in crickets. To determine if *L. equestris* can physiologically perceive bat body odor, we employed a GC–EAD system following established methods (Barbosa-Cornelio et al., 2019; Ma et al., 2025) to test the electrophysiological sensitivity of *L. equestris* antennae to bat-derived VOCs. The system consisted of an Agilent 7820A gas chromatograph (Agilent Technologies, USA) and EAD equipment (Syntech, Germany). The EAD setup comprised an IDAC 4 signal recording controller, an MP-15 micromanipulator with high-impedance PRG-3 electrodes.

The tips and bases of cricket antennae were trimmed and mounted on a PRG-3 electrode using an MP-15 micromanipulator. After achieving a stable baseline, odor samples were separated on an Agilent 7820A gas chromatograph. A 4 μL aliquot was injected in splitless mode onto an HP-5 column (30 m × 0.32 mm × 0.25 μm). We selected the HP!Il5 column because its non!Ilpolar stationary phase achieves excellent chromatographic separation of the non!Ilpolar to weakly polar hydrocarbons that dominate bat body odor, which is essential for subsequent detection. The GC oven temperature was programmed as follows: held at 50 °C for 5 min, increased at 10 °C/min to 200 °C (held 1 min), then raised at 20 °C/min to 250 °C (held 5 min). Nitrogen carrier gas flow was 2.0 mL/min. Injector and detector temperatures were 250 °C and 300 °C, respectively. The GC effluent was split at a 1:2 ratio using a Y-shaped splitter. Two-thirds of the effluent was directed to a flame ionization detector (FID) for compound detection, and one-third was transferred through a heated transfer line into a humidified airstream (400 mL/min) before being delivered to the cricket antenna. Antennal signals were acquired using an IDAC-4 interface and GcEad software (v. 1.2.5; Syntech, Germany), allowing simultaneous recording of FID and EAD responses. Each chromatographic peak in the FID trace had a corresponding EAD trace showing the depolarization of the antennal potential.

A total of eight antennae from individual crickets were tested. Five antennae were exposed to bat odor extracts, and three were exposed to empty-chamber controls (one antenna per injection). The empty-chamber controls were used to account for potential background signals from the experimental system and to provide a baseline for comparison with bat odor extracts. Snout secretions were not included in the GC–EAD analyses because the present study focused on bat whole-body odor as the ecologically relevant airborne odor source, and sufficient secretion samples for standardized volatile extraction were not available for electrophysiological testing. A compound was considered EAD-active if its GC peak coincided with an antennal depolarization >0.1□mV, and this response was reproducible in at least three individual antennae. Active compounds were identified by aligning the retention times and peak profiles of EAD-active signals with those from parallel GC–MS analyses.

### Chemical profiling and source analysis of *S. kuhlii* body odor

To characterize the major volatile constituents of *S. kuhlii* body odor, we analyzed the collected VOC extracts by GC–MS using an Agilent 6850-5975 mass spectrometer operated in electron ionization (EI) mode at 70 eV. The GC was equipped with the same HP-5 column used for GC–EAD, with helium as the carrier gas at a constant flow of 1 mL/min. A 1 μL aliquot of each sample was injected in splitless mode (0.75-min splitless period) at an injector temperature of 250 °C.

The oven temperature program matched that used for GC–EAD. Compounds were tentatively identified by comparison of their mass spectra with the NIST 2017 library (Supplementary Table 4; Supplementary Figure 2–7). The identities of these compounds were confirmed by matching both retention times and mass spectra to those of authentic standards, including injections of a mixture containing 100 ng of each compound. However, co-injection with authentic standards could not be performed because the total extract volume from each sample was insufficient for additional runs after the primary GC-MS and GC-EAD analyses, owing to the limited amount of biological material available.

To investigate potential biological sources and characterize the volatile chemical profiles of bat body odor, we performed a broadly targeted volatile metabolomic analysis using headspace solid–phase microextraction coupled with GC–MS (HS–SPME–GC–MS) on hair, pararhinal gland (snout) secretions, and feces from nine bats (four females, five males). These bats were collected and maintained as described above. Upon capture, bats were placed in clean stainless-steel cages and kept in groups consistent with their natural social associations during the brief interval prior to immediate odor sampling. Hair samples (10 mg per individual) were clipped from dorsal and ventral regions. Snout secretions were collected using sterile cotton swabs (CS15-005, Shenzhen SihuaBo Technology Co., Ltd., China), with two blank swabs as controls. These blank swab controls were included to account for potential volatile contamination from ambient air or the swab material itself (Supplementary Table 5). Fecal samples (100 mg per individual) were collected as in the dietary analysis. All samples were stored in 2-ml Eppendorf tubes at −80 °C until analysis.

Samples were thawed on ice, transferred to headspace vials, and spiked with 10 µL of an internal standard (50 µg/mL). Volatiles were extracted by HS–SPME: after incubation at 60□°C for 5□min, a 120□μm DVB/CWR/PDMS fiber was exposed to the headspace for 15□min for adsorption, then thermally desorbed at 250□°C for 5□min in the GC injector. Analysis was performed on an Agilent 7890B 7000D GC–MS system equipped with a DB 5MS capillary column (30□m□×□0.25□mm□×□0.25□μm), with helium at 1.2□mL/min. The oven temperature program was: 40□°C for 3.5□min, then increased at 10□°C□/min to 100□°C, 7□°C/min to 180□°C, and 25□°C/min to 280□°C (hold 5□min). The MS operated in EI mode at 70□eV, with ion source, quadrupole, and interface temperatures set at 230□°C, 150□°C, and 280□°C, respectively. Data were acquired in selected ion monitoring (SIM) mode. Each compound was identified by matching the retention time and the presence of one quantitative and two to three qualitative ions to those of a standard reference (Yuan et al., 2022). Data were integrated, corrected, and processed using MassHunter software (B.08.00; Agilent), and metabolites were qualified and quantified against an in-house database (Supplementary Table 5). Retention times for non-target compounds are not fully disclosed due to the proprietary policy of the commercial analytical service provider.

To identify the biological source of the characteristic body odor, we compared the VOC profiles from hair, feces, and snout secretions. Volatiles were classified into mutually exclusive chemical classes for comparison; note that “Hydrocarbons” here refers only to aliphatic hydrocarbons, and “Nitrogen compounds” excludes amines. To objectively assess the overall compositional similarity among hair, feces, snout secretions, and body odor, we performed a principal component analysis (PCA) based on binary (presence/absence) VOC data.

### Identifying avoidance-eliciting odor compounds

Chemical profiling of *S. kuhlii* body odor by GC–MS identified six volatile compounds: 2,2-dimethylheptane, limonene, undecane, 2,6,7-trimethyldecane, pentadecane, and hexadecane, two of which elicited EAD responses (Figure 2C). Because GC–EAD provides a physiological readout of antennal sensitivity but may not fully capture behavioral relevance across different concentrations or odor contexts, we additionally tested antennal responses to commercially available standards of compounds identified by GC–MS. We selected four compounds (limonene, undecane, pentadecane, and hexadecane) and evaluated antennal responses using electroantennography (EAG) (Supplementary Table 6). Each compound was diluted to 10% (v/v) in hexane for screening, resulting in a final limonene concentration of 5.87 × 10^−7^ mol/µL. Hexane alone served as the solvent control. Stimuli were prepared by applying 10 μL of a test solution to a filter-paper strip (10 mm × 30 mm), which was then placed inside a glass Pasteur pipette. The pipette tip was positioned in a humidified airstream (1500 mL/min) directed at the antenna through a stainless-steel tube (7 mm inner diameter, 1 cm from the antenna). Cricket antennae were prepared as for GC–EAD. For each antenna, a hexane control was first presented to establish baseline antennal activity. For the initial screening, limonene, undecane, pentadecane, and hexadecane were diluted to 10% (v/v) in hexane and delivered individually in a randomized order. To assess dose-dependent responses, limonene was further tested at five concentrations (0.001%, 0.01%, 0.1%, 1%, and 10%, v/v in *n*-hexane), following the concentration gradient used in a previous study (Tang et al., 2024). Following the initial hexane control, the five limonene concentrations were tested in a randomized order across trials. Each stimulus lasted 0.5 s, with an inter-stimulus interval of 1 min to allow full recovery of antennal responses. EAG responses were recorded from eight antennae for the initial screening and from six antennae for the dose–response assay, with each antenna obtained from a different individual, and analyzed using EagPro software (version 2.0; Syntech, Germany).

To examine the behavioral response of *L. equestris* to limonene, a two-choice assay was conducted using 10% (v/v) limonene (5.87 × 10^−7^ mol/µL) as the odor source. A 10 μL aliquot was applied to a filter-paper strip (10 mm × 30 mm) inside a glass chamber. A total of 38 *L. equestris* individuals were tested under the same experimental conditions as in the bat-odor assay.

To ecologically validate that a single odor compound can elicit avoidance, we tested the effect of 10% (v/v) limonene (5.87 × 10^−7^ mol/µL) on the calling activity of *L. equestris* within the bats’ foraging habitat. Over nine nights between 24 June to 16 July 2024, we established four sampling plots (5 m × 5 m) in a linear array at 100-m intervals within the known foraging range of *S. kuhlii*, at the same location where crickets were originally captured (Supplementary Figure 8). Each night, two plots were randomly assigned to the limonene treatment and two to hexane control. To assess avoidance, we monitored changes in calling activity of *L. equestris* as a proxy, using passive acoustic monitoring during their peak nocturnal activity. At each plot, an Audiomoth (v 1.2.0, Open Acoustic Devices, UK) was positioned 0.5 m above ground with the microphone facing downward. Audio was recorded in 55-s files separated by 5-s intervals (1-min duty cycle) at a 384 kHz sampling rate (medium gain). A 40-min pre-exposure recording started at 19:50. Between 20:30 and 20:40, 1 mL of 10% limonene (treatment) or hexane (control) was evenly sprayed over the 25 m^2^ area centered on the recorder. A 40-min post-exposure recording started at 20:50. Experimental nights were separated by at least one day, and rainy conditions were avoided.

During acoustic preprocessing, we excluded plot–night datasets that failed quality-control criteria: no cricket calls detected, >20% of 1-min files with zero calls, or total call count <10,000. This quality-control procedure yielded a final sample of 15 datasets: 8 from the limonene-treated (experimental) plots and 7 from the hexane-treated (control) plots. In total, 1,200 recordings (80 files/plot/night) were analyzed. Using Avisoft SASLab Pro (v 5.2.07, Avisoft Bioacoustics, Germany), recordings were band-pass filtered (3–6.5 kHz) to match the calling frequency of *L. equestris* (dominant frequency ∼5 kHz, range 4.5–5.5 kHz). This frequency band does not overlap with sympatric cricket species, and *L. equestris* is the dominant local cricket (see Results), allowing reliable identification. Calls were identified via spectrogram matching using 10 templates of *L. equestris* calls. A “call” was defined as a train of successive sound elements (syllables). Typical calls of *L. equestris* last 60–200 ms (Supplementary Figure 9), with inter-syllable intervals < 10 ms within a call, and >60 ms between consecutive calls from the same individual. The number of calls per minute was quantified for each plot and recording period. Our assays were designed to test physiological detectability and functional sufficiency, rather than to establish concentration thresholds or mimic natural emissions exactly.

### Data analysis

All analyses were performed in R v4.3.2 unless specified otherwise. For the two-choice olfactory assay, we used a chi-square goodness-of-fit test to compare the number of crickets exhibiting “avoiding” versus “not avoiding” behavior. Effect sizes are reported as the phi coefficient (φ) with 95% confidence intervals (estimated via the delta method). For chemical profiling, we performed multivariate analysis on VOC data. Hierarchical clustering with heatmap visualization was conducted in Python (v3.9) using the *pandas*, *seaborn*, and *matplotlib* libraries, based on data normalized to a 0–1 range, Euclidean distance, and average linkage. Principal component analysis (PCA) based on a binary (presence/absence) matrix was performed using the *vegan* package in *R* to compare profiles from hair, feces, snout secretions, and bat body odor. For the EAG experiment, EAG responses were analyzed using repeated-measures ANOVA (sphericity checked with the *rstatix* package), followed by Bonferroni-corrected paired *t*-tests. Effect sizes are reported as partial eta-squared (η²*p*) for the ANOVA and Hedges’ *g* for the *t*-tests. For field calling activity, a generalized linear mixed model (GLMM) was fitted to calls per minute using the *glmmTMB* package. Model selection compared zero-inflated and standard negative binomial distributions using AIC, and evaluated alternative random effects structures and fixed effects using AICc. The final model included a Group × Phase interaction as a fixed effect, with a random intercept for the plot-by-date combination to account for spatial and temporal non-independence. Model diagnostics were performed with the *DHARMa* package, including tests for residual uniformity, overdispersion, and zero-inflation.

### Animal welfare and ethics

The bats used in this study were carefully handled and maintained under controlled conditions (28 °C, 60% relative humidity) with *ad libitum* access to food and water. Bats were housed socially in cages (50 cm × 50 cm × 50 cm). No bat mortality occurred during the study, and all bats were released at their original roost sites upon completion of experiments. Crickets were kept at densities ≤ 5 individuals per container to minimize aggression. Crickets not used in EAG assays were returned to their collection sites. All procedures involving live animals were approved by the Science and Technology Ethics Committee of Northeast Normal University, Changchun, China (permit ID: NENU-2022-0308).

## Supporting information

Supplementary Figures and Tables_revised

## Acknowledgements

We are grateful to Jiqian Li, Yinli Hu, and Maojun Zhong for their advice on diet and behavioral analysis, and to Weiwei Wang and Pengji Li for their assistance with the field experiments.

## Funding

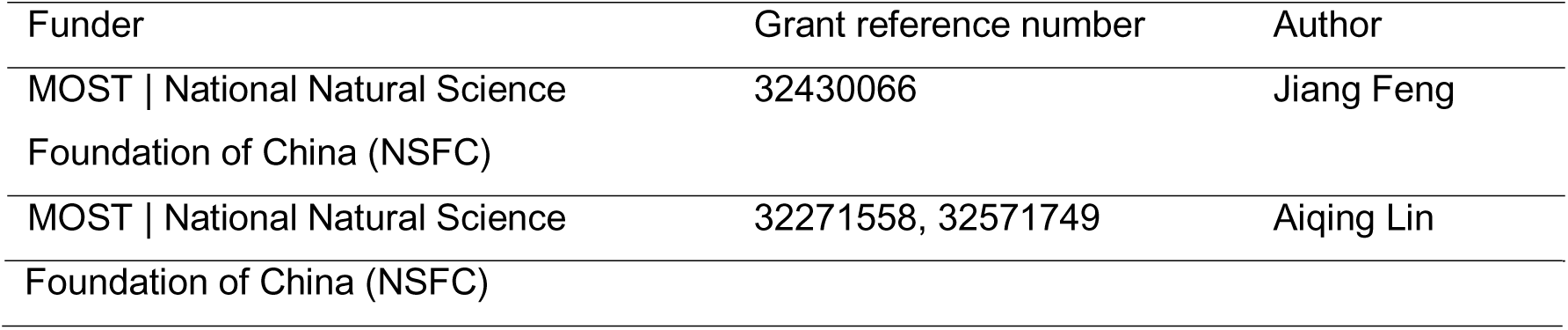

## Author contributions

Y.L., A.L. and J.F. designed research; Y.L., W.Z., J.W., and H.X. performed research; Y.L. and A.L. analyzed data and wrote the paper.

## Data availability

The processed volatile compound data (chemical identities, relative abundances, retention indices, and mass spectral details) for *S. kuhlii* body odor, feces, hair, and secretions are provided in Supplementary Table 4 and 5. Representative GC–MS chromatograms and reference mass spectra for key compounds are provided in Supplementary Figures 2–7. The raw behavioral, electrophysiological, and ecological datasets have been deposited in Figshare and will be publicly accessible upon manuscript acceptance. During peer review, these datasets can be accessed confidentially via the private link: https://figshare.com/s/e6af2463a2d36062bf0e

## Supplemental Information

**Supplementary Figure 1. GC–EAD responses of *L. equestris* antennae to *S. kuhlii* body odor.** Upper panel shows the flame ionization detection (FID) chromatogram of *S. kuhlii* body odor extract, with peaks 1–6 corresponding to compounds identified by GC–MS. The five EAD traces show antennal responses from five individual *L. equestris* crickets exposed to the same bat body odor extract (*n* = 5 crickets). Asterisks indicate EAD active peaks that elicited antennal depolarizations. Lower panel shows the FID chromatogram and EAD traces from odor-free control samples, which did not elicit antennal responses in three tested crickets (*n* = 3 crickets).

**Supplementary Figure 2. GC–MS identification of compound 1 from the body odor of *Scotophilus kuhlii.*** (A) Experimental EI mass spectrum of compound 1 identified from the volatile profiles of *S. kuhlii*. (B) Standard EI mass spectrum of 2,2-dimethylheptane for comparison. The horizontal axis represents the mass-to-charge ratio (m/z), and the vertical axis represents the relative intensity (%).

**Supplementary Figure 3. GC–MS identification of compound 2 from the body odor of *Scotophilus kuhlii.*** (A) Experimental EI mass spectrum of compound 2 identified from the volatile profiles of *S. kuhlii*. (B) Standard EI mass spectrum of limonene for comparison. The horizontal axis represents the mass-to-charge ratio (*m/z*), and the vertical axis represents the relative intensity (%).

**Supplementary Figure 4. GC–MS identification of compound 3 from the body odor of *Scotophilus kuhlii.*** (A) Experimental EI mass spectrum of compound 3 identified from the volatile profiles of *S. kuhlii*. (B) Standard EI mass spectrum of undecane for comparison. The horizontal axis represents the mass-to-charge ratio (*m/z*), and the vertical axis represents the relative intensity (%).

**Supplementary Figure 5. GC–MS identification of compound 4 from the body odor of *Scotophilus kuhlii.*** (A) Experimental EI mass spectrum of compound 4 identified from the volatile profiles of *S. kuhlii*. (B) Standard EI mass spectrum of 2,6,7-trimethyldecane for comparison. The horizontal axis represents the mass-to-charge ratio (*m/z*), and the vertical axis represents the relative intensity (%).

**Supplementary Figure 6. GC–MS identification of compound 5 from the body odor of *Scotophilus kuhlii.*** (A) Experimental EI mass spectrum of compound 5 identified from the volatile profiles of *S. kuhlii*. (B) Standard EI mass spectrum of pentadecane for comparison. The horizontal axis represents the mass-to-charge ratio (*m/z*), and the vertical axis represents the relative intensity (%).

**Supplementary Figure 7. GC–MS identification of compound 6 from the body odor of *Scotophilus kuhlii.*** (A) Experimental EI mass spectrum of compound 6 identified from the volatile profiles of *S. kuhlii*. (B) Standard EI mass spectrum of hexadecane for comparison. The horizontal axis represents the mass-to-charge ratio (*m/z*), and the vertical axis represents the relative intensity (%).

**Supplementary Figure 8. Study site and field setup.** (A) Satellite image of the roosting habitat of the insectivorous bat *Scotophilus kuhlii*, indicating the locations of bat roosting trees and experimental quadrats. (B) A representative quadrat established within the natural habitat of the cricket *Loxoblemmus equestris* for field experiment. (C) An Audiomoth acoustic recorder deployed inside a quadrat.

**Supplementary Figure 9. Waveform and spectrogram characterizing the calling song of *Loxoblemmus equestris*.** The upper panel depicts the oscillogram of a male calling song, while the lower panel presents the corresponding spectrogram, plotting frequency (kHz) against time (s).

**Supplementary Table 1. Composition of Gryllidae (crickets) in the insect prey community at a foraging site of *Scotophilus kuhlii*.** Species identified from grassland samples collected between July and September 2022 are listed with their corresponding order, family, and number of individuals.

**Supplementary Table 2. Annotation information for terpenoid compounds detected in VOC samples from *Scotophilus kuhlii*.** Key parameters used for volatile organic compound (VOC) annotation are summarized, including the NIST retention index (RI), quantitative ion, qualitative ion(s), and molecular weight (MW, Da). Compound identification was based on matching retention time and the presence of one quantitative ion and two to three qualitative ions with standard references, followed by qualification and quantification against an in-house database.

**Supplementary Table 3. Generalized linear mixed models examining the effects of limonene (experimental) versus hexane (control) treatments on call activity in the cricket *Loxoblemmus equestris*.** The model was fitted using a zero-inflated negative binomial distribution via the *glmmTMB* package in R. Fixed effects included the four-level treatment–phase combination (Control–Pre-exposure, Control–Post-exposure, Experimental–Pre-exposure, Experimental–Post-exposure); random effects comprised a random intercept for plot-night (plot:date). Results are presented twice with different reference categories: first with ‘Control–Pre-exposure’, then with ‘Experimental–Pre-exposure’. Both presentations derive from the same fitted model, yielding two intercept estimates but identical coefficients.

**Supplementary Table 4. Volatile compounds identified from the body odor of *Scotophilus kuhlii* by GC–MS analysis.** Compounds are listed with their CAS registry numbers, retention times, corrected peak areas, relative abundances, mass spectral match factors, reverse match factors, verification database, and electroantennographic detection (EAD) activity.

**Supplementary Table 5. Volatile compounds identified from bat feces, hair, snout secretions, and blank swab controls using HS-SPME-GC–MS analysis.** Compounds are listed with their CAS registry numbers, retention times, retention indices, quantitative and qualitative ions, molecular weights, molecular formulas, chemical classes, and relative signal intensities detected in different sample types.

**Supplementary Table 6. Synthetic chemical standards and solvents used in the study.**

