## Supplementary Figures and Tables_revised for "Beyond Acoustic Cues: Olfactory-Mediated Avoidance of Bats by Crickets"

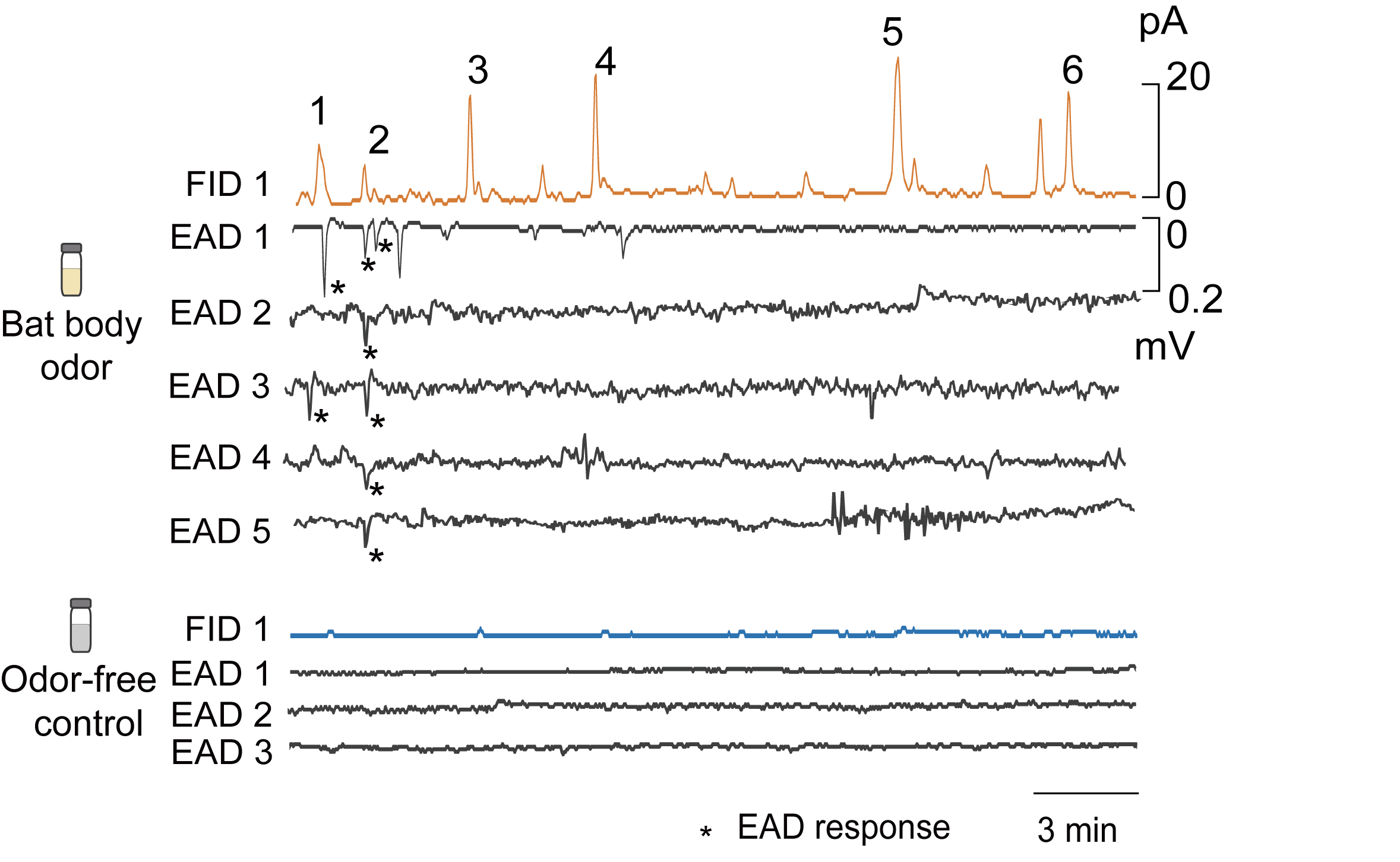

**Supplementary Figure 1.** **GC–EAD responses of *L. equestris* antennae to *S. kuhlii* body odor.** Upper panel shows the flame ionization detection (FID) chromatogram of *S. kuhlii* body odor extract, with peaks 1–6 corresponding to compounds identified by GC–MS. The five EAD traces show antennal responses from five individual *L. equestris* crickets exposed to the same bat body odor extract (*n* = 5 crickets). Asterisks indicate EAD active peaks that elicited antennal depolarizations. Lower panel shows the FID chromatogram and EAD traces from odor-free control samples, which did not elicit antennal responses in three tested crickets (*n* = 3 crickets).

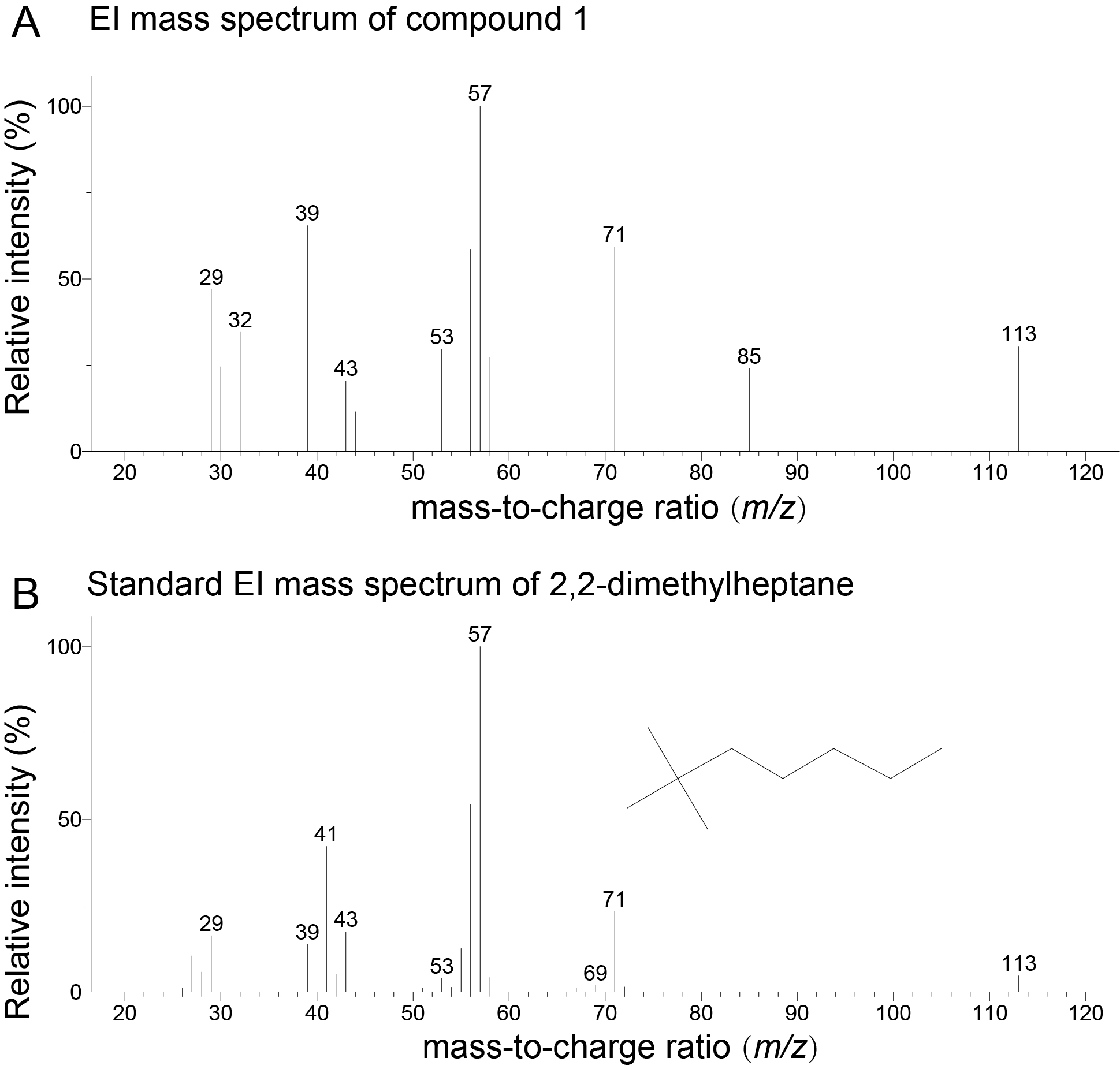

**Supplementary Figure 2. GC–MS identification of compound 1 from the body odor of *Scotophilus kuhlii.*** (A) Experimental EI mass spectrum of compound 1 identified from the volatile profiles of *S. kuhlii*. (B) Standard EI mass spectrum of 2,2-dimethylheptane for comparison. The horizontal axis represents the mass-to-charge ratio (m/z), and the vertical axis represents the relative intensity (%).

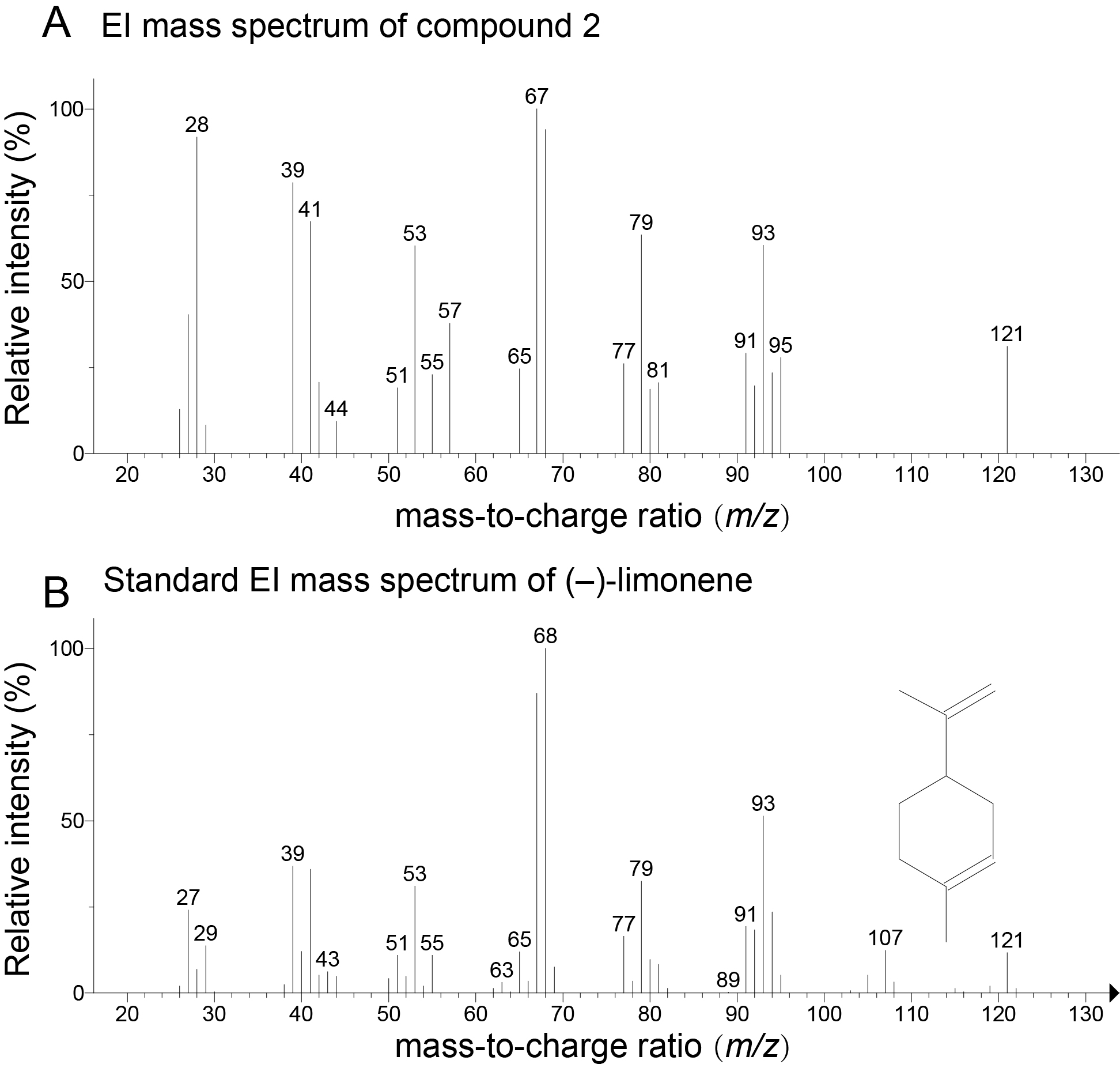

**Supplementary Figure 3. GC–MS identification of compound 2 from the body odor of *Scotophilus kuhlii.*** (A) Experimental EI mass spectrum of compound 2 identified from the volatile profiles of *S. kuhlii*. (B) Standard EI mass spectrum of limonene for comparison. The horizontal axis represents the mass-to-charge ratio (*m/z*), and the vertical axis represents the relative intensity (%).

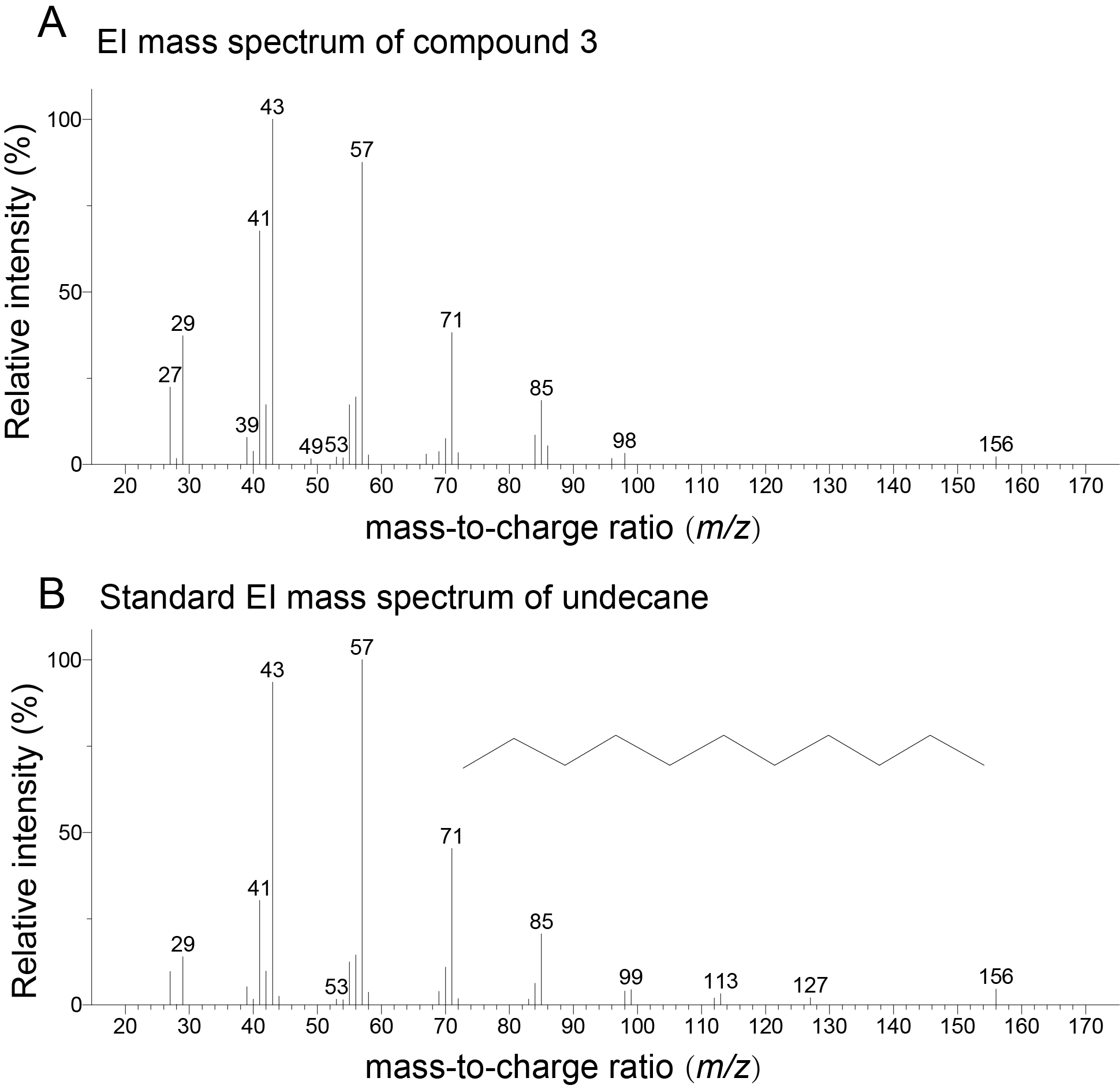

**Supplementary Figure 4. GC–MS identification of compound 3 from the body odor of *Scotophilus kuhlii.*** (A) Experimental EI mass spectrum of compound 3 identified from the volatile profiles of *S. kuhlii*. (B) Standard EI mass spectrum of undecane for comparison. The horizontal axis represents the mass-to-charge ratio (*m/z*), and the vertical axis represents the relative intensity (%).

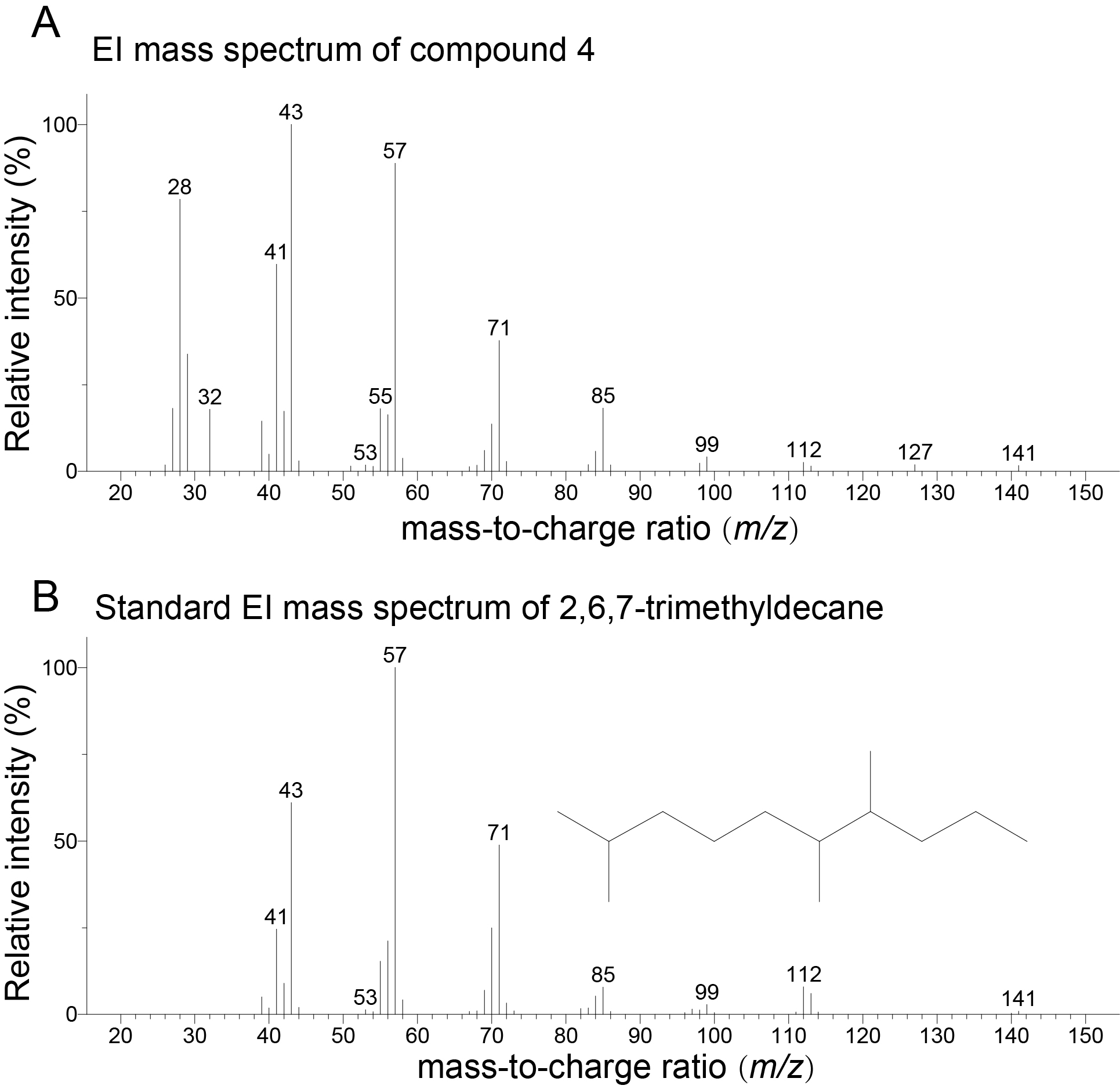

**Supplementary Figure 5. GC–MS identification of compound 4 from the body odor of *Scotophilus kuhlii.*** (A) Experimental EI mass spectrum of compound 4 identified from the volatile profiles of *S. kuhlii*. (B) Standard EI mass spectrum of 2,6,7-trimethyldecane for comparison. The horizontal axis represents the mass-to-charge ratio (*m/z*), and the vertical axis represents the relative intensity (%).

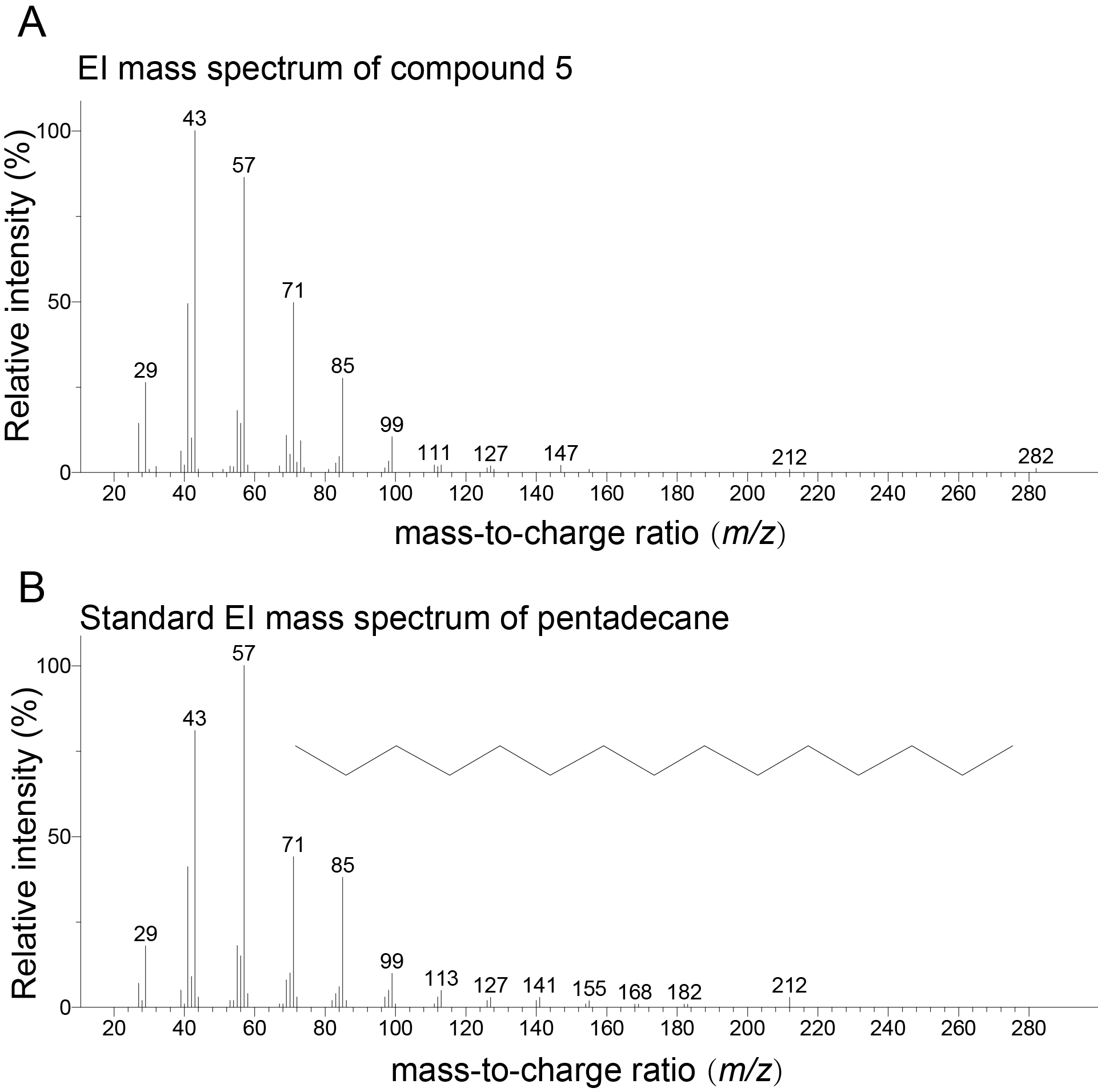

**Supplementary Figure 6. GC–MS identification of compound 5 from the body odor of *Scotophilus kuhlii.*** (A) Experimental EI mass spectrum of compound 5 identified from the volatile profiles of *S. kuhlii*. (B) Standard EI mass spectrum of pentadecane for comparison. The horizontal axis represents the mass-to-charge ratio (*m/z*), and the vertical axis represents the relative intensity (%).

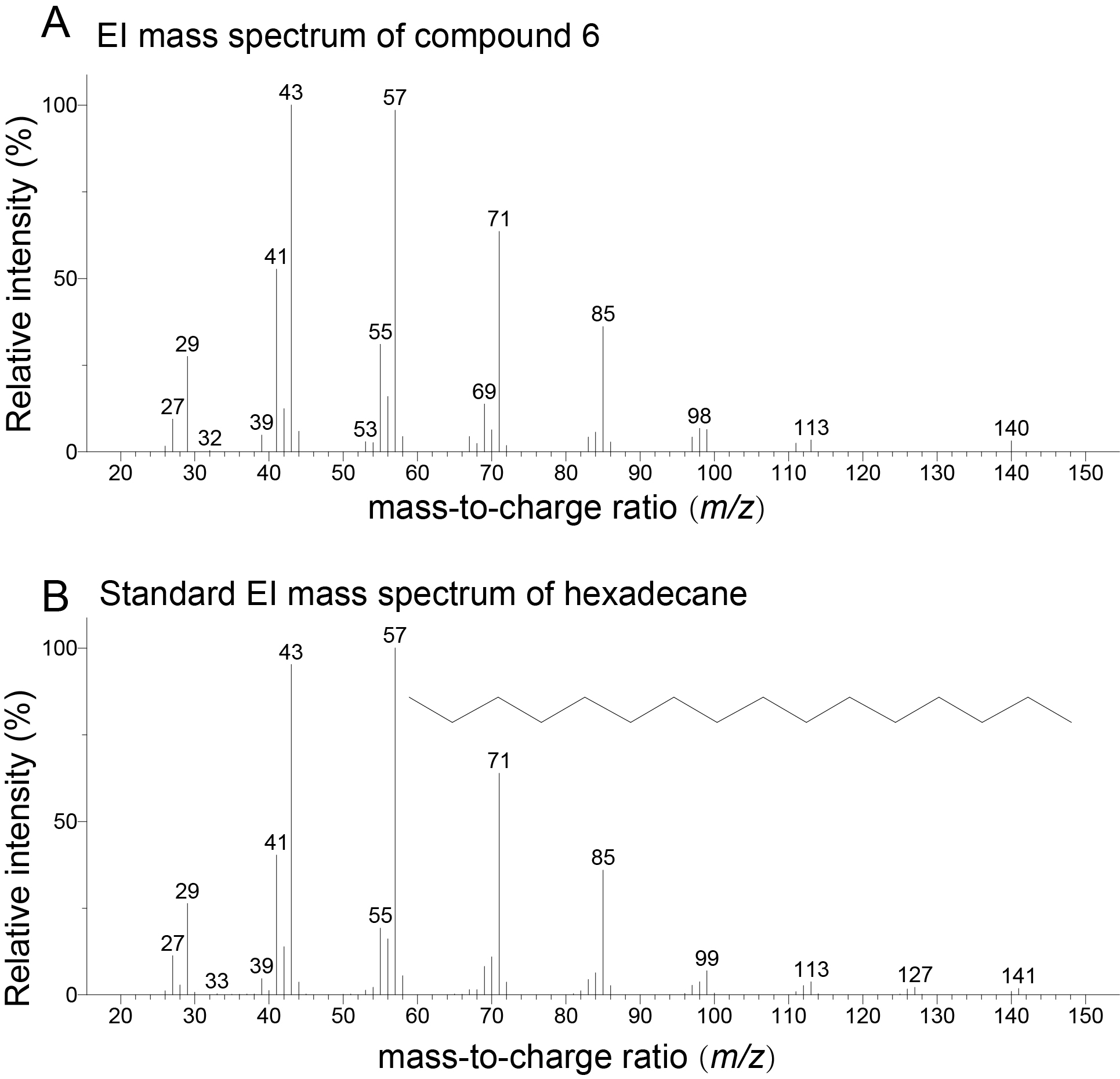
**Supplementary Figure 7. GC–MS identification of compound 6 from the body odor of *Scotophilus kuhlii.*** (A) Experimental EI mass spectrum of compound 6 identified from the volatile profiles of *S. kuhlii*. (B) Standard EI mass spectrum of hexadecane for comparison. The horizontal axis represents the mass-to-charge ratio (*m/z*), and the vertical axis represents the relative intensity (%).

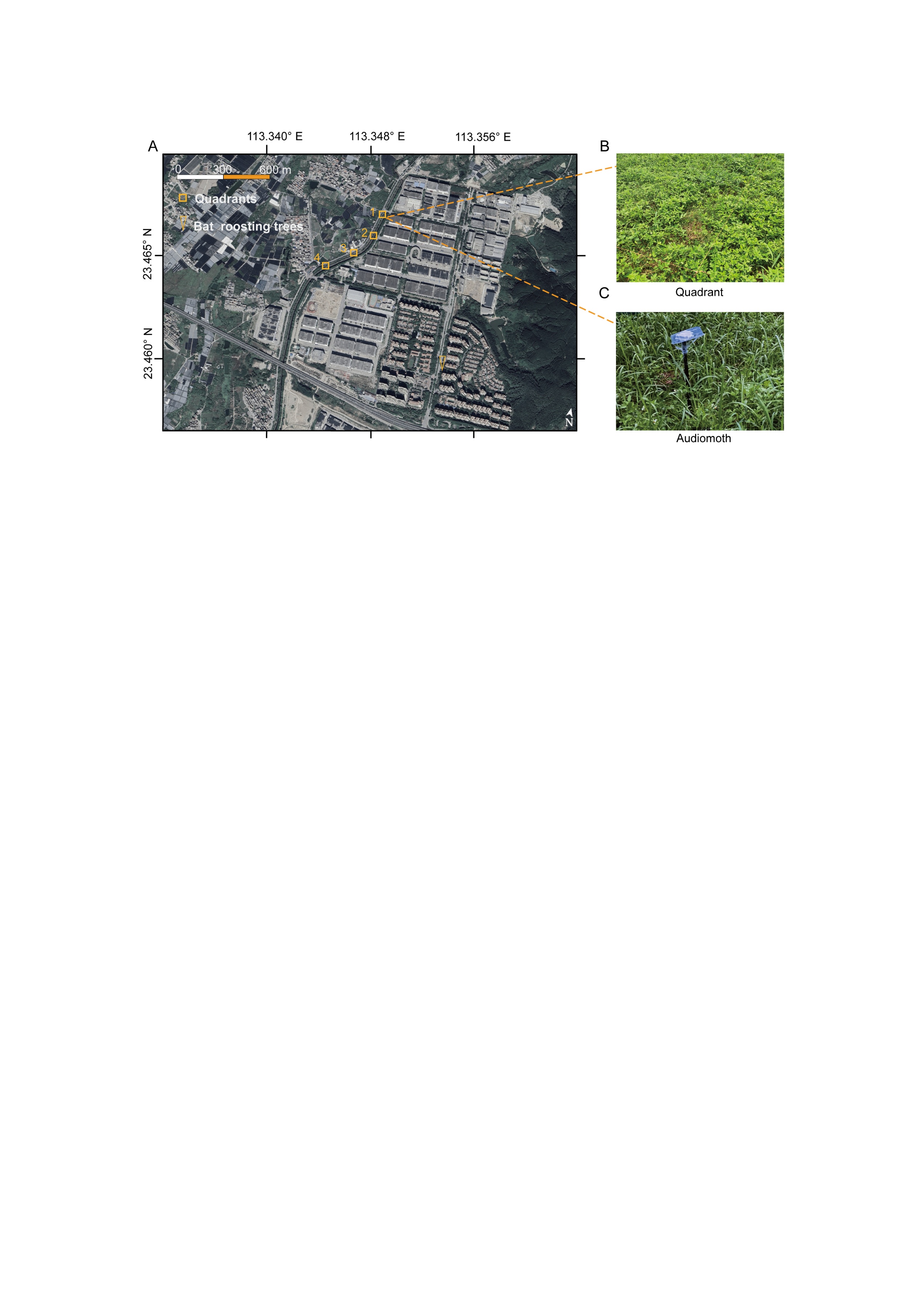

**Supplementary Figure 8. Study site and field setup.** (A) Satellite image of the roosting habitat of the insectivorous bat *Scotophilus kuhlii*, indicating the locations of bat roosting trees and experimental quadrats. (B) A representative quadrat established within the natural habitat of the cricket *Loxoblemmus equestris* for field experiment. (C) An Audiomoth acoustic recorder deployed inside a quadrat.

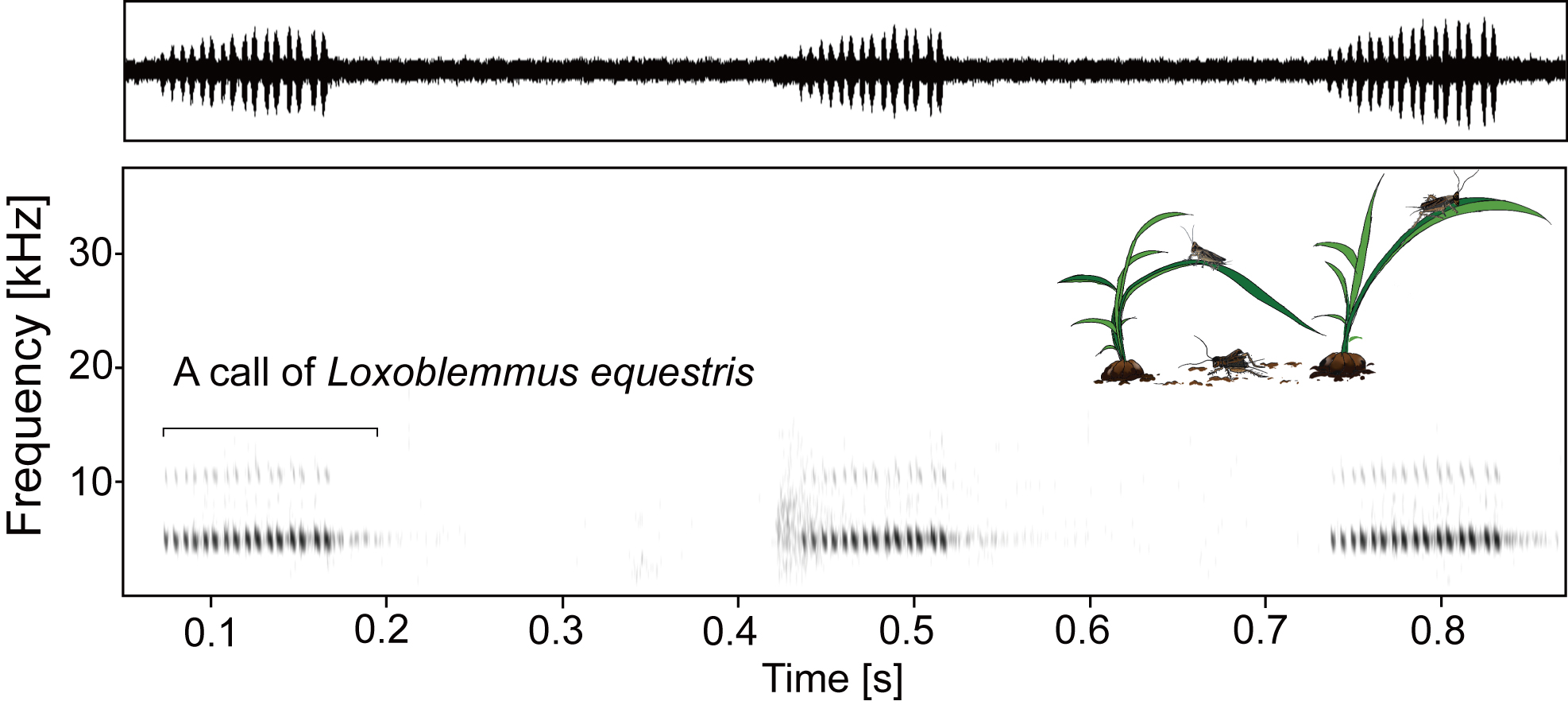

**Supplementary Figure 9. Waveform and spectrogram characterizing the calling song of *Loxoblemmus equestris*.** The upper panel depicts the oscillogram of a male calling song, while the lower panel presents the corresponding spectrogram, plotting frequency (kHz) against time (s).

**Supplementary Table 1. Composition of Gryllidae (crickets) in the insect prey community at a foraging site of *Scotophilus kuhlii*.** Species identified from grassland samples collected between July and September 2022 are listed with their corresponding order, family, and number of individuals.

| Insect species | Order | Family | Number |
| --- | --- | --- | --- |
| *Loxoblemmus equestris* | Orthoptera | Gryllidae | 70 |
| *Polionemobius taprobanensis* | Orthoptera | Gryllidae | 1 |
| *Svercacheta siamensis* | Orthoptera | Gryllidae | 5 |
| *Amusurgus genji* | Orthoptera | Gryllidae | 1 |
| *Modicogryllus consobrinus* | Orthoptera | Gryllidae | 1 |
| *Teleogryllus emma* | Orthoptera | Gryllidae | 2 |

**Supplementary Table 2. Annotation information for terpenoid compounds detected in VOC samples from *Scotophilus kuhlii*.** Key parameters used for volatile organic compound (VOC) annotation are summarized, including the NIST retention index (RI), quantitative ion, qualitative ion(s), and molecular weight (MW, Da). Compound identification was based on matching retention time and the presence of one quantitative ion and two to three qualitative ions with standard references, followed by qualification and quantification against an in-house database.

| **Compound** | **NIST RI** | **Quant. ion** | **Qual. ion** | **MW (Da)** |
| --- | --- | --- | --- | --- |
| Limonene | 1023 | 93 | 136 | 136.125 |
| ar-Curcumene | 1524 | 119 | 132 | 202.172 |
| α-Phellandrene | 969 | 93 | 91 | 136.125 |
| δ-Elemene | 1377 | 121 | 93 | 204.188 |
| trans-Farnesol | 1710 | 41 | 69 | 222.198 |
| Terpinolene | 1052 | 93 | 121 | 136.125 |
| Nerolidol | 1564 | 93 | 69 | 222.198 |
| Phytol | 2045 | 71 | 43 | 296.308 |
| δ-Cadinene | 1469 | 161 | 134 | 204.188 |
| Pulegone | 1212 | 152 | 81 | 152.12 |
| Cedrol | 1543 | 95 | 150 | 222.198 |
| β-Elemene | 1398 | 81 | 68 | 204.188 |
| Safranal | 1186 | 107 | 91 | 150.104 |
| 7-Octylidene-bicycloheptane | 1522 | 135 | 93 | 206.203 |

**Supplementary Table 3. Generalized linear mixed models examining the effects of limonene (experimental) versus hexane (control) treatments on call activity in the cricket *Loxoblemmus equestris*.** The model was fitted using a zero-inflated negative binomial distribution via the *glmmTMB* package in R. Fixed effects included the four-level treatment–phase combination (Control–Pre-exposure, Control–Post-exposure, Experimental–Pre-exposure, Experimental–Post-exposure); random effects comprised a random intercept for plot-night (plot:date). Results are presented twice with different reference categories: first with ‘Control–Pre-exposure’, then with ‘Experimental–Pre-exposure’. Both presentations derive from the same fitted model, yielding two intercept estimates but identical coefficients.

| Term | Estimate | SE | 95% CI | *Z* | *P* |
| --- | --- | --- | --- | --- | --- |
| Control–Pre-exposure (Intercept) | 5.30 | 0.18 | [4.94, 5.66] | 29.02 | <0.001 |
| Control–Post-exposure | 0.40 | 0.06 | [0.28, 0.51] | 6.68 | <0.001 |
| Experimental–Post-exposure | -0.59 | 0.25 | [-1.08, -0.10] | -2.35 | 0.019 |
| Experimental–Pre-exposure | 0.10 | 0.25 | [-0.39, 0.59] | 0.38 | 0.720 |
| Experimental–Pre-exposure (Intercept) | 5.40 | 0.17 | [5.06, 5.73] | 31.59 | <0.001 |
| Experimental–Post-exposure | -0.68 | 0.06 | [-0.80, -0.57] | -11.90 | <0.001 |
| Control–Post-exposure | 0.30 | 0.25 | [-0.19, 0.79] | 1.21 | 0.228 |
| Control–Pre-exposure | -0.10 | 0.25 | [-0.59, 0.39] | -0.38 | 0.702 |
